# Identification of druggable host dependency factors shared by multiple SARS-CoV-2 variants of concern

**DOI:** 10.1101/2023.01.09.523209

**Authors:** Ilaria Frasson, Linda Diamante, Manuela Zangrossi, Elena Carbognin, Anna Dalla Pietà, Alessandro Penna, Antonio Rosato, Ranieri Verin, Filippo Torrigiani, Cristiano Salata, Lorenzo Vaccaro, Davide Cacchiarelli, Sara N. Richter, Marco Montagner, Graziano Martello

**Author notes:** These authors contributed equally (Ilaria Frasson, Linda Diamante, Manuela Zangrossi). These authors jointly supervised this work (Sara N. Richter, Marco Montagner, Graziano Martello).

## Abstract

The high mutation rate of SARS-CoV-2 leads to emergence of several variants, some of which are resistant to vaccines and drugs targeting viral elements. Targeting host dependency factors – cell proteins required for viral replication - would help avoid resistance. However, whether different SARS-CoV-2 variants induce conserved cell responses and exploit the same core host factors is still unclear.

We compared three variants of concern and observed that the host transcriptional response was conserved, differing only in kinetics and magnitude. By CRISPR screening we identified the host genes required for infection by each variant: most of the identified genes were shared by multiple variants, both in lung and colon cells. We validated our hits with small molecules and repurposed FDA-approved drugs. All drugs were highly effective against all tested variants, including delta and omicron, new variants that emerged during the study. Mechanistically, we identified ROS production as a pivotal step in early virus propagation. Antioxidant drugs, such as N-acetyl cysteine (NAC), were effective against all variants both in human lung cells, and in a humanised mouse model. Our study supports the use of available antioxidant drugs, such as NAC, as a general and effective anti-COVID-19 approach.

## INTRODUCTION

The continuous emergence of multiple variants is an intrinsic feature of several viruses and of the pandemic caused b y SARS-CoV-2. This poses important hurdles to the development of prophylactic approaches as new variants partially overcome the immunity generated by COVID-19 vaccines(1). Recently, two first-generation antivirals have been approved. However, targeting viral proteins will also eventually lead to selection of resistant variants(2). Inhibiting host-factors might prove a better strategy to avoid resistance, given that host-factors are obviously not under selective pressure to favour viral propagation. Besides the differences in transmissibility and clinical severity, we currently do not know whether different variants share the same life cycle or if they exploit different molecular routes in the host cell: in the first case new variant-independent druggable targets could be envisaged. Unfortunately, direct comparison of the requirements and life cycles among the different variants is still missing due to the multitude of model systems (cell lines, organoids and transgenic animals) and variants used in the different studies (3).

Previous genetic screenings for host-factors involved in SARS-CoV-2 infection highlighted the role of several cellular proteins (4–8), but the overlap between different studies remains very low. We still do not know whether this poor overlap is due to actual heterogeneity in the cellular proteins exploited by the different variants for their propagations, or rather to technical differences. Here, we analysed the transcriptional responses of human lung cells to three SARS-CoV-2 variants that sequentially overtook each other during the pandemic, namely Wuhan, D614G and Alpha variants, and performed a high-stringency CRISPR-based genetic loss-of-function screen to identify host-factors necessary for the infection by these variants. We identified 525 genes, most of which shared by two or more variants. Gene ontology analysis showed an enrichment of terms related to mitochondrial organisation and oxidative stress. We performed individual validation of CRISPR gene candidates by RNA interference and selected proteins against which currently available drugs are available for potential COVID-19 repurposing. We provide evidence that genetic and chemical inhibition of RIPK4, SLC7A11 and MASTL leads to strong inhibition of virus-induced cytotoxicity. Further, we validated our findings on two additional variants (Delta and Omicron), which emerged during our study. We focused on SLC7A11 and show that interfering with viral-induced ROS accumulation hinders viral replication suggesting that these targets and compounds might be effective against current and future variants of SARS-CoV-2.

## RESULTS

To investigate SARS-CoV-2 biology and its interactions with the human host cell, we selected three major SARS-CoV-2 variants of concern that emerged and spread worldwide up to late 2020. The Wuhan variant is the original strain that emerged in Wuhan, China at the end of 2019; the Wuhan D614G variant emerged soon after, in early 2020, and contains the stated mutation which has been maintained in all the later variants and is thought to enhance viral replication; the Alpha variant (B.1.1.7), first detected in December 2020, contains several mutations in the spike protein that mark it out from the original Wuhan strain and make it more contagious (up to 50% more transmissible) and associated with increased disease severity (Supplementary Table 1)(9). Since the first and main target of infection of all SARS-CoV-2 variants is the respiratory tract, where severe pneumonia can develop, we employed Calu-3 cells, a human lung epithelial cell line that is both susceptible (i.e. it expresses both ACE2 receptor and TMPRSS2 cofactor on the cell membrane) and permissive to SARS-CoV-2 infection (10). This is the only human lung cell line that allows efficient SARS-CoV-2 replication with production of high amounts of viral progeny resulting in cell death (11, 12). Other human lung cell lines are susceptible but only marginally permissive (13). Alternative cell lines were made susceptible with ACE-2 overexpression via plasmid transfection (10, 14). In the two latter cases no detectable cytopathic effect upon viral release could be measured.

We first compared replication of Wuhan, D614G and Alpha strains in Calu-3 cells infected at multiplicity of infection (MOI) of 0.1(15) by measuring: 1) the amount of intracellular viral RNA and 2) the amount of new infectious viral particles produced at different hours post infection (h.p.i.)(16, 17) (Supplementary Fig. 1). At both 24 and 48 h.p.i., the Wuhan virus produced the lowest number of viral transcripts, and the highest amounts of new infective particles; vice versa, Alpha was very efficient in viral transcript production, while it generated the lowest amount of new infective particles. The D614G variant displayed an intermediate behaviour, with high RNA transcription rates and copious viral progeny generation. Both parameters highly increased from 24 to 48 h.p.i. for Wuhan and D614G, while they remained constant for Alpha. We concluded that the 3 variants display differences in their replication rates and viral transcription(15–18) (Supplementary Fig. 1). All 3 variants caused cell death of Calu-3 within 48 h.p.i.(19–21), but the Alpha variant was more rapid and led to complete cell death by 24 h.p.i., in agreement with the more rapid viral transcript production.

We next set out to study both viral and cellular transcriptome changes upon infection by RNA sequencing. To maximise the number of infected cells and thus avoid confounding data from uninfected cells, we increased MOI to 1. To assess the impact during the first cycle of viral replication only, we tested shorter times post infection: 6, 9, 12 and 24 h.p.i.. At 24 h.p.i. Alpha led to complete death of infected cells, so it could not be analysed. We first observed that viral transcripts were readily detectable after 6 hours in D614G and Alpha and that, in general, Alpha showed the most pronounced transcriptional response, followed by D614G and Wuhan (Fig. 1a), as also confirmed by quantitative Real-Time PCR (qPCR, Supplementary Fig. 2a). These results are in line with those obtained at lower MOI (Supplementary Fig. 1).

**Figure 1.**
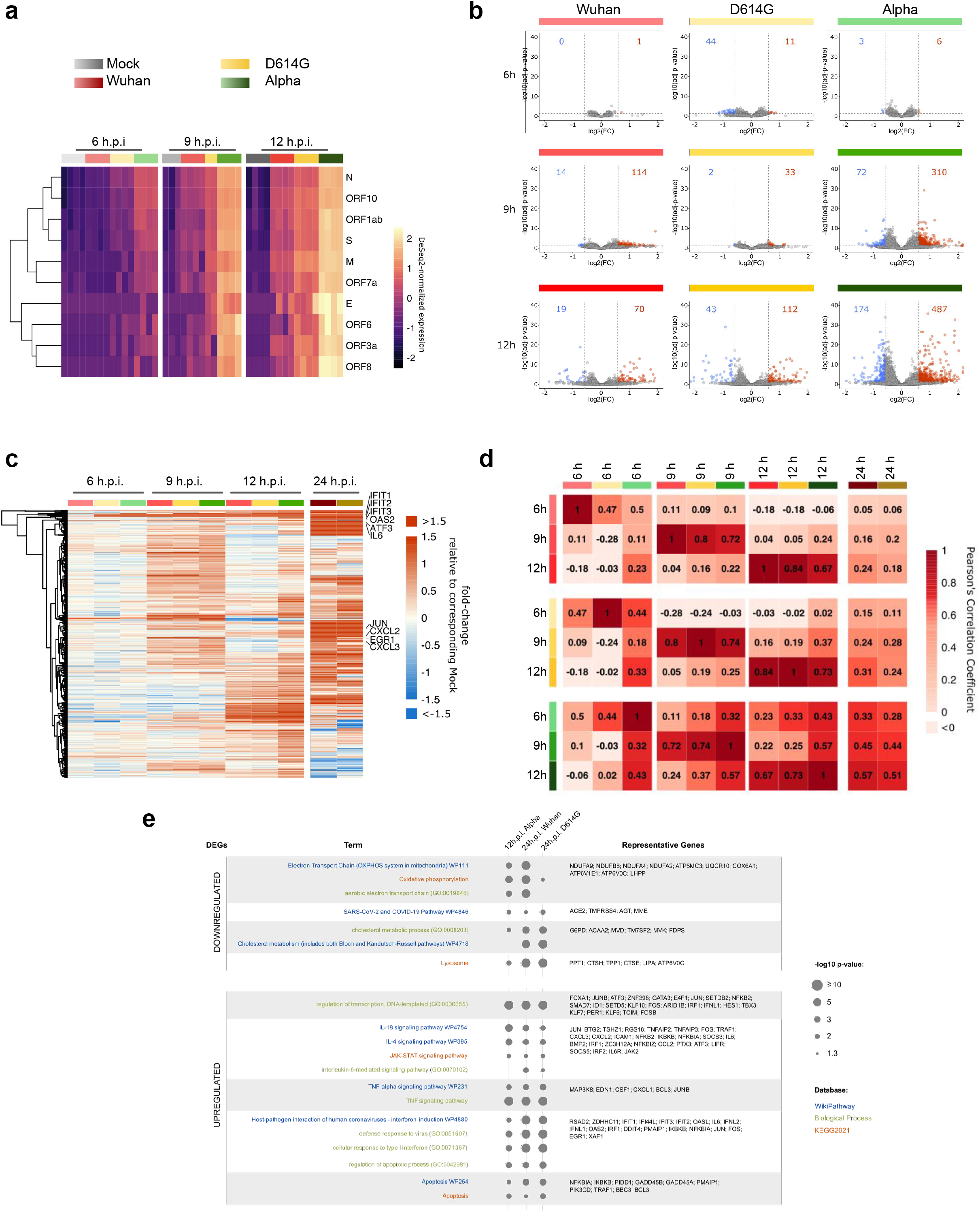
Different SARS-CoV-2 variants induce highly similar transcriptional responses. **a**, Heatmap of viral transcripts in uninfected cells (mock) or cells infected with the indicated variants. Expression is shown as row-scaled Z-scores. **b**, Volcano plots showing differentially expressed genes (DEGS, log(FC)>0.59 or <-0.59, adjusted p-value <0.05) between uninfected and infected cells at the indicated time points. The number of upregulated and downregulated DEGs are indicated in red and blue, respectively. **c**, Heatmap of the fold-change of upregulated DEGs within 12 h.p.i.. **d**, Correlation matrix displaying Pearson’s correlation among the indicated samples. **e**, Gene enrichment analysis on DEGs identified in samples infected with Alpha variant for 12 h.

To test whether the observed differences in viral transcription kinetics translated into different cellular responses, we concurrently analysed the expression levels of cellular transcripts and found a high number of differentially expressed genes (DEGs) starting from 9 h.p.i. (Fig. 1b). The kinetics of cellular gene expression followed a pattern similar to that of viral transcription, i.e. the magnitude of variation was the highest with Alpha, followed by D614G and Wuhan variants (Fig. 1b). Most of the modulated genes were similarly regulated among the different variants, although Alpha induced stronger and quicker responses (Fig. 1b-c). At 24 h.p.i. Wuhan and D614G led to gene expression patterns similar to those from Alpha at 12 h.p.i. (Fig. 1c). These data confirm that Alpha displays quicker infection kinetics, but also indicate that the modulated genes, even though affected at different times, are shared among the tested variants. Several of the modulated genes have been previously reported to be induced by SARS-CoV-2 infection *in vitro* and in patients(22, 23) (e.g. IL6, IFIT1/2/3, OAS2, CXCL2, ATF3 and EGR1). We confirmed the differences in kinetics and magnitude of the transcriptional response in host genes by qPCR (Supplementary Fig. 2b).

To further investigate the similarities in transcriptional responses we calculated the correlation coefficient among the different variants during infection (Fig. 1d). A value of 1 indicates that a gene has the same magnitude of expression in the two variants that are compared. At 6 h.p.i., we observed weak correlations, in line with the absence of a robust transcriptional response (Fig. 1c). However, from 9-12 h.p.i. we obtained increased correlation coefficients among the genes induced by the three variants (Fig. 1d), indicating convergence towards similar gene expression profiles. Comparison against publicly available transcriptomic data revealed highly significant overlap of DEGs with other studies based on different coronavirus variants and species (Supplementary Fig. 3). To identify cellular functions underlying the expression of different gene modules, we performed gene enrichment analysis (Fig. 1e). Downregulated genes describe a cellular context with decreased mitochondrial respiration, decreased cholesterol synthesis and reduced expression of ACE2 and TMPRSS4, key mediators of SARS-CoV-2 entry. On the contrary, processes such as gene transcription, Interferon, Interleukin, JAK-STAT, TNF signalling pathways and response to human coronaviruses, were highly activated, in line with previous studies(20–24) (Fig. 1e). Finally, we noticed an increased expression of apoptosis-related genes, in line with cell death observed upon Alpha infection.

Our transcriptional analyses revealed high similarities among the 3 variants both at the virus and cell level, with differences mainly at the temporal level; however this did not inform us on the host dependency factors shared by the different variants. Previous studies identified cellular genes required for infection of SARS-CoV-2(4–8) via CRISPR-based genetic loss-of-function screens that detect cell genes, the expression of which is essential for viral replication. Each study used a different combination of virus variants and host cells. Overlap among the genes identified in the different studies was very limited (Supplementary Fig. 4), as also reported by Baggen and colleagues(8). We thus resolved to perform a systematic comparison of the 3 variants in Calu-3 cells under identical experimental conditions (Fig. 2a and Supplementary Fig. 5). Calu-3 cells, besides deriving from one of the most relevant tissue targets of SARS-CoV-2 infection and morbidity, are compatible with knock-out screening due to their hypotriploid karyotype(3). We performed high stringency CRISPR-based loss-of-function screening by infecting cells with SARS-CoV-2 at MOI 3, at which we observed complete Calu-3 cell death within 48 h.p.i. The rationale was that only knocked-out mutant cells where viral infection was abrogated would survive. To this end, Cas9-expressing Calu-3 cells were stably transduced with a genome-wide library of guide RNA (gRNA) to direct Cas9 and knock-out a single gene per cell. The integrated gRNA worked both as a guide for Cas9 and as a barcode for the identification of the targeted locus. Non-transduced cells were used as control to check for complete virus-induced cell death. Genomic DNA from uninfected cells and from surviving clones was purified and gRNA identified by NGS. We used high coverage (500X, in triplicates for each variant) and, to reduce the risk of false positives and off-targets, we excluded gRNAs that targeted genes expressed at no or very low levels in our transcriptional analysis (Fig. 1b). We then set up the “gRNA score” as the number of biological replicates in which a given gRNA was found enriched in infected over uninfected cells (see Methods). The gRNA score was calculated within each SARS-CoV-2 variant (ranging from 0 to 3 replicates) or combining all variants (ranging from 0 to 9 replicates).

**Figure 2.**
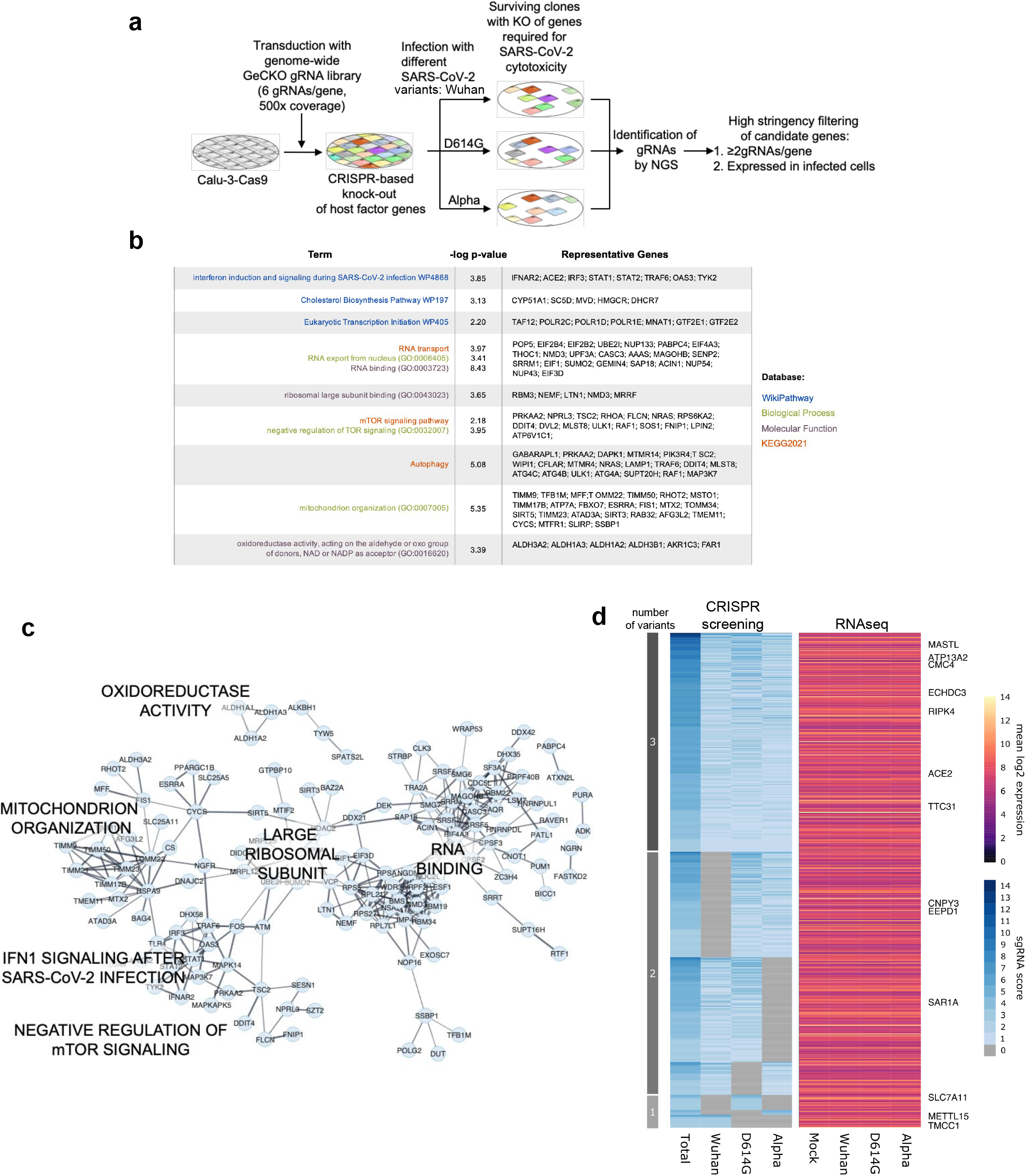
High stringency CRISPR-based loss-of-function screening provides biological insights on different SARS-CoV-2 variants. **a**, Schematics of the screening strategy. **b**, Gene enrichment analysis on hits identified by the CRISPR screening in at least one variant. **c**, Protein network of candidate hits emerging from the screening. Edges show connections based on experiments, co-expression, text mining, databases, cooccurrence, neighbourhood, fusion (any) and edges width is proportional to the strength of the interaction (mapping type “continuous” based on “Stringdb score” value). **d**, Left, heatmap of the gRNA scores for genes shared by 3, 2 or specific for 1 variant. Right, mean expression levels at 12 h.p.i. of candidate genes, shown as log2 normalised expression.

We first focused on genes with gRNA score >1, i.e. genes whose knockout led to cell survival upon infection with at least one SARS-CoV-2 variant (see Methods and Supplementary Table 2) and we filtered out genes not robustly expressed in Calu-3. Gene enrichment analysis identified terms previously associated with SARS-CoV-2 infection, such as host-pathogen interaction with human coronaviruses, interferon induction and cholesterol biosynthesis, indicating that the screening procedure was successful(4, 24, 25). Interestingly, this analysis also identified ribosomal organisation, oxidoreductase activity and mitochondrial organisation, as top regulated processes (Fig. 2b, c), in line with our transcriptomic analysis results (Fig. 1e). We then applied more stringent filters, and considered only hit genes with gRNA score >2, which were identified by at least 2 independent gRNAs. We identified 525 genes, 44.2% were shared among the 3 variants, 49.1% among 2 variants and only 6.7% were found in only 1 variant (Fig. 2d). Importantly, the expression levels of the majority (95.3%) of candidates were not changed by infection with any variants under consideration (Fig. 2d, right). These results indicate that different SARS-CoV-2 variants exploit a highly shared set of host factors that are constitutively expressed and not affected by the infection.

Next, we performed secondary validation by transient short interfering RNA (siRNA)-mediated knockdown. We chose siRNAs as a loss-of-function strategy because it is totally unrelated to CRISPR, and it is used in the clinic(26). The hits identified in our genetic screening could in principle act at different stages of the infection cycle, from receptor binding to virus assembly and cell exit: to monitor all viral steps in a single assay, we tested the end-point release of infective viral progeny from infected cells. In the primary CRISPR-based screening only a small number (35/525) of genes resulted specific to only 1 variant (Fig. 2d), potentially indicating that different variants rely on the same host proteins for infection, or simply reflecting technical limitations of the screening procedure. To clarify this aspect, we chose genes shared by 1, 2 or 3 variants, with robust and stable expression during infection, and challenged each silenced hit with the three viral variants. We used ACE2-, TMPRSS2-, TMPRSS4- and STAT2-silenced samples as positive controls of viral infection inhibition, given their known role in viral entry and Interferon response(27). Efficient siRNA-mediated knockdown of each hit (Supplementary Fig. 6a) led to a dramatic decrease in the viral titre of all 3 variants (Fig. 3a). Silencing of the hits reduced production of new viral particles to a level comparable to that achieved with the positive controls. These data corroborate the accuracy and strength of our CRISPR-based screening.

**Figure 3.**
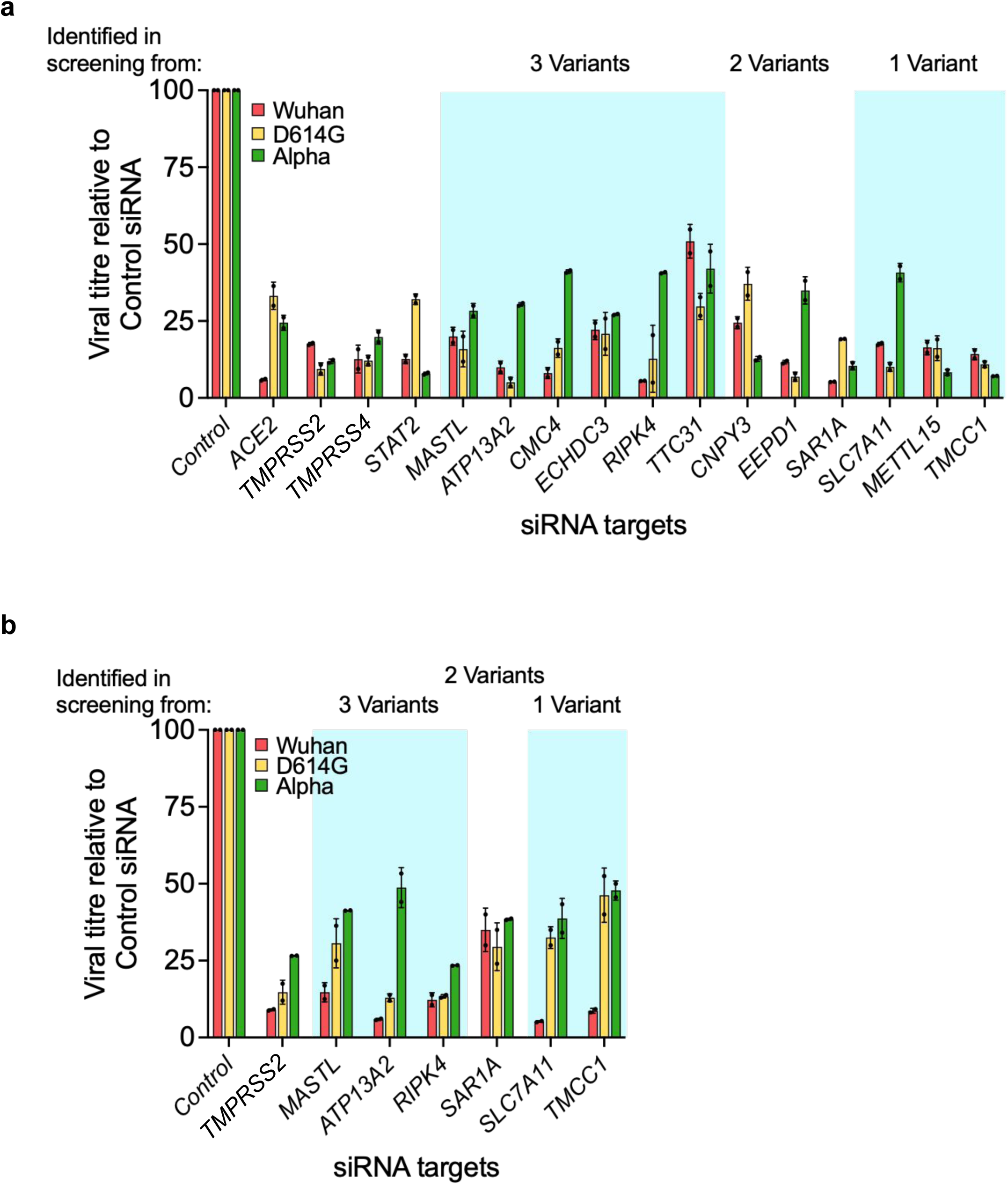
Inhibition of the infection of different SARS-CoV-2 strains by siRNA-mediated downregulation of screening-retrieved human genes. Calu-3 (**a**) and Caco-2 cells (**b**) treated with non-targeting (CTR) or indicated siRNAs and infected with SARS-CoV-2 (used variants are shown, MOI 0.1). Infectious SARS-CoV-2 particles in the supernatant were assessed by plaque reduction assay. Bars represent the mean of two independent replicates (± s.d.). “3, 2, 1 variants” indicate the number of variants against which the shown hit genes were found.

SARS-CoV-2 was reported to infect to a lesser extent also cells from the gastrointestinal tract, inducing gastrointestinal manifestations in the human host with consequent viral shedding in stools(28, 29): Thus, we further validated by siRNA knock-down a subselection of hits in Caco-2 cell line, which derives from colonic epithelium and is susceptible and permissive to SARS-CoV-2 infection(30) (Supplementary Fig. 6b). As observed in lung cells, knockdown of hit genes significantly reduced the viral titre of all 3 variants in colon cells, indicating conservation of the hits across different tissue types (Fig. 3b).

In light of providing hits with faster clinical translatability, we then searched for FDA approved drugs and drug-like small molecules targeting our hits and tested them for antiviral activity. We focused on RIPK4, SLC7A11 and MASTL, 3 proteins that have never been previously associated with SARS-CoV-2 infection. Receptor-interacting serine/threonine-protein kinase 4 (RIPK4) interacts with protein kinase C (PKC) β and PKCδ, and regulates keratinocyte differentiation, cutaneous inflammation, and cutaneous wound repair(31). RIPK4 is inhibited by Tamatinib and Vandetanib(32, 33). SLC7A11, also known as xCT, is the major subunit of the cystine/glutamate antiporter that has been extensively studied for its role in the regulation of the cell redox state, and thus in cell homeostasis and the pathophysiology of several diseases (34–36). In viruses, SLC7A11 has been shown to mediate entry and post-entry events of the Kaposi Sarcoma-Associated Herpes sVirus(37–39). This antiporter is inhibited by Sulfasalazine and Imidazole ketone Erastin (IKE)^18,19^. Microtubule-associated serine/threonine kinase-like (MASTL) regulates mitosis and meiosis and it is considered a promising anticancer target. A novel compound, named MKI-1, was recently described as a specific inhibitor of MASTL(42).

We first tested all compounds for their cytotoxic activity on Calu-3 cells (Supplementary Table 3) and calculated CC_50_ values. Most compounds displayed cytotoxicity in the micromolar range (Tamatinib, Vandetanib and IKE), MKI-1 was cytotoxic in the high nanomolar range, whereas Sulfasalazine displayed no cytotoxicity at all tested concentrations. Next, the antiviral activity of all compounds was tested at concentrations at which no cytotoxicity on host cells was detected to avoid confounding effects. All compounds were tested on the three described virus variants, and the occurrence and spreading of the Delta variant by the time we reached this phase of the study, prompted us to validate all antiviral candidates also against it (43). All tested compounds showed strong, dose dependent and selective antiviral effect, measured as inhibition of viral progeny production at 48 h.p.i. and expressed as IC_50_ value and selectivity index (SI), i.e. the ratio between CC_50_ and IC_50_ (Fig. 4 a-e). Tamatinib, Vandetanib, IKE and MKI-1 were all highly active in the nanomolar range and inhibited virus production by 85-93% at the highest tested concentration. IKE and Vandetanib both displayed an excellent average SI (around 100, Fig. 4a-d). Sulfasalazine induced a strong antiviral effect in the low micromolar range (IC_50_ 5.7 *μ*M), and retained an acceptable SI (> 20) due to its negligible cytotoxicity (Fig. 4e). Notably, all tested compounds were also active against the Delta variant, further supporting the “variant-wide” relevance of our hit genes. Importantly, 3 out of 5 tested compounds are FDA approved drugs, and hence amenable for drug-repurposing studies. RIPK4, SLC7A11 and MASTL hence emerge as bona fide therapeutic targets for multiple SARS-CoV-2 variants.

**Figure 4.**
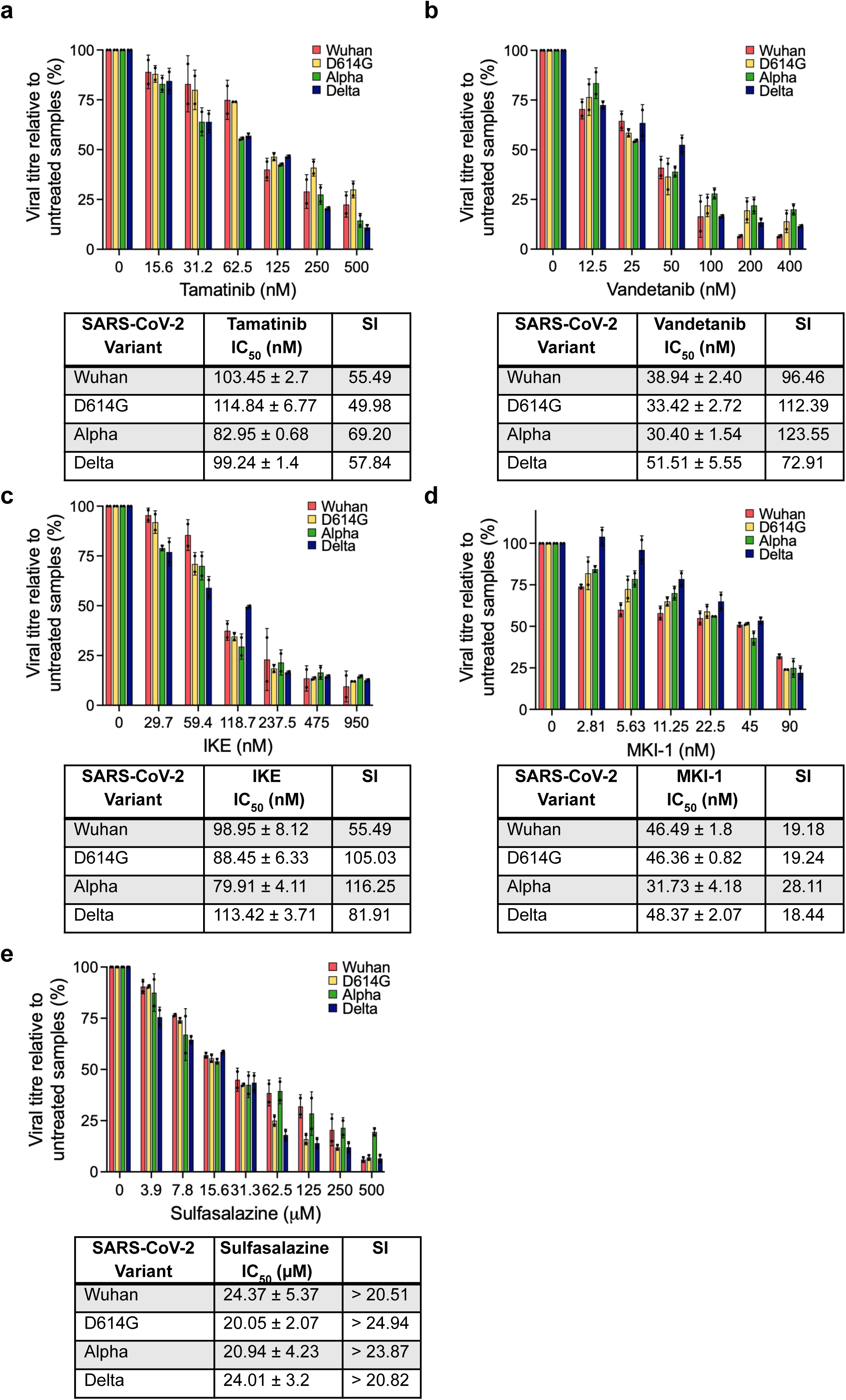
Assessment of the antiviral activity in human lung cells of the compounds hindering the cellular targets. Calu-3 cells were pretreated for 24 h with the indicated compounds and infected with SARS-CoV-2 (strains Wuhan, D614G, Alpha and Delta, MOI 0.1). Two days post infection, cell medium was subjected to plaque reduction assay and the viral titre was calculated and expressed as PFU/ml. Data are mean ± s.d. of n=2 biological replicates. Each condition was tested in triplicate per replicate.

Our genome-wide screening identified mitochondrial and oxidoreductive activities as crucial components of the cellular response to SARS-CoV-2 infections, suggesting a potential role for ROS. Indeed, SLC7A11 is a key regulator of the cell redox state, thus we investigated if the antiviral mechanism of SLC7A11 inhibitors relied on intracellular ROS modulation. To this end we focused on IKE as SLC7A11 inhibitor and excluded Sulfasalazine, because several independent studies recently reported higher risk of deaths for COVID-19 in patients treated with Sulfasalazine. This effect is opposite to our in vitro findings and possibly related to SLC7A11-independent inhibition of type I interferon (IFN) production in vivo(44).

We assessed intracellular ROS levels upon 24 h treatment of Calu-3 cells with IKE and observed a biphasic trend, with an initial increase at early time points followed by a substantial decrease (Fig. 5a). This is compatible with the secondary activation of compensatory mechanisms, already reported after long term inhibition of SLC7A11 with erastin and IKE(45, 46). We then tested intracellular ROS levels upon SARS-CoV-2 infection at different times post-infection and within the first replication cycle(17, 47). In our conditions, we observed a sharp increase in ROS levels at 2 h.p.i., while at longer times ROS returned to basal levels (Fig. 5b), in line with the reported intracellular inflammation due to Spike protein binding to the ACE2 receptor and the subsequent expression of non-structural viral proteins(48, 49). In contrast, pretreatment (24 h) with IKE prior to viral infection(50–54), reduced basal ROS levels in cells (Fig. 5b, Time 0). Upon infection, ROS levels remained low for 30 minutes in IKE treated cells (Fig. 5b, Time 0.5-2). At longer times of infection, ROS levels were comparable to those of untreated and uninfected cells, in both IKE-treated and untreated infected cells. Thus, IKE pretreatment in infected cells prevented virus-induced ROS burst (up to 2 h.p.i.).

**Figure 5.**
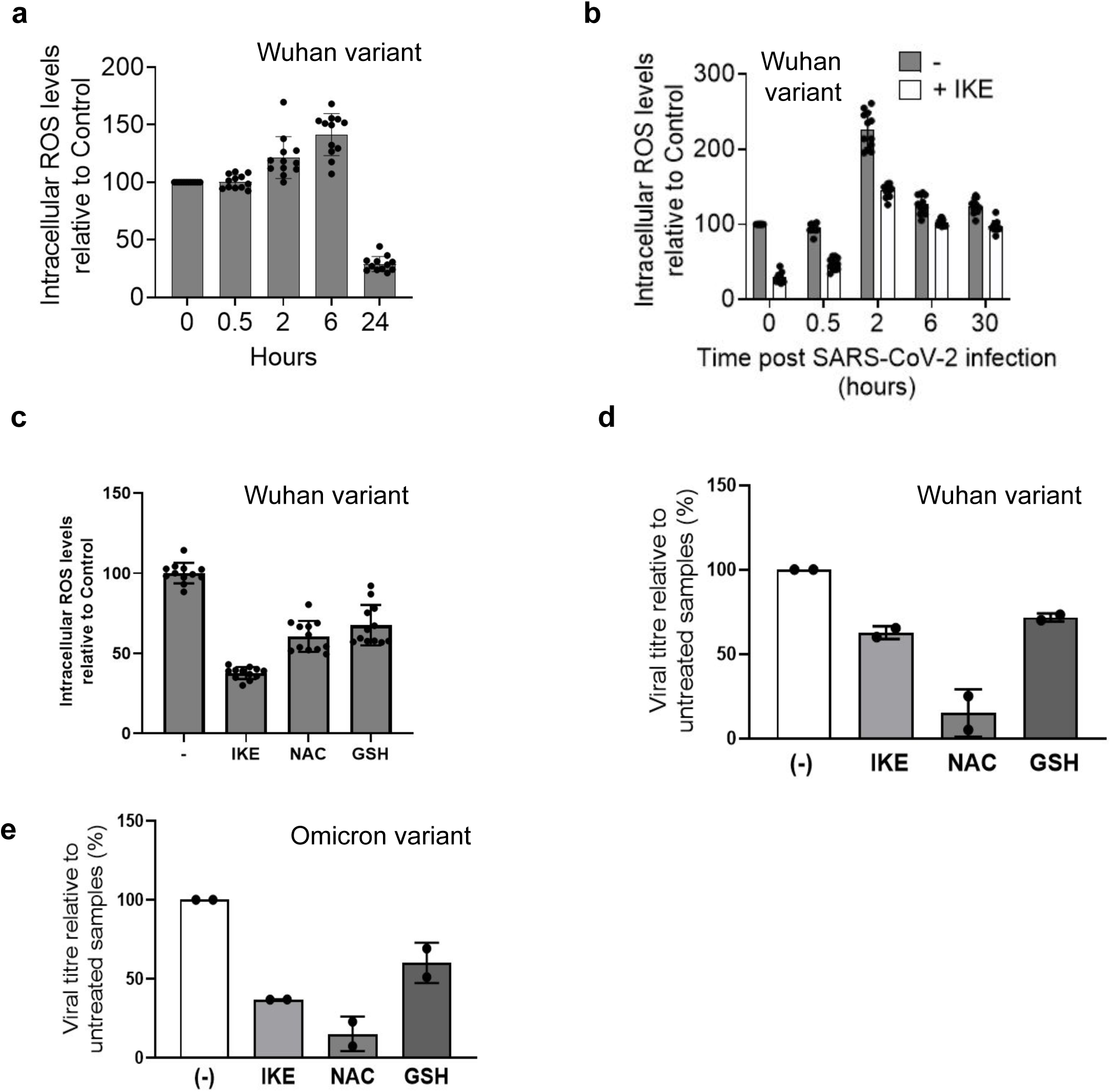
Decreasing ROS levels impairs viral replication. **a** Intracellular ROS levels were measured in Calu-3 cells at different times after IKE administration (950 nM). IKE prompted a modest ROS increase from 2 to 6 h post administration, whereas longer times of exposure, induced ROS depletion. **b** Intracellular ROS levels were measured at different times post SARS-CoV-2 infection in Calu-3 cells, in the absence or presence of IKE (950 nM). In the latter case, cells were also pretreated with IKE 24 hours prior to infection. Data were normalized to the untreated control of each tested time point. The viral titre was assessed by PRA, each condition was assayed in = 3 samples. Data are mean ± s.d. of n = 2 biological replicates. **c** Intracellular ROS levels were measured in Calu-3 cells upon IKE (950 nM) or GSH (300 μM) administration (24 h treatment) and compared to untreated cells (-) as control. Both IKE and GSH induced ROS depletion in Calu-3 cells. IKE treatment diminished ROS levels up to the 70% with respect to the control, whereas GSH administration reduced intracellular ROS up to the 40%. Intracellular ROS levels were determined by H2dcfda assay, each conditions was evaluated in sextuplicate. Single values ± s.d. of n=2 biological replicates are shown. **d** Viral titre (Wuhan variant) upon 24 h pre-treatment of Calu-3 cells with IKE (950 nM) or GSH (300 μM) and testing after a single cycle of replication, i.e. 30 hpi. **e** Viral titre (Omicron variant) upon 24 h pre-treatment of Calu-3 cells with IKE (950 nM) or GSH (300 μM) and testing after a single cycle of replication, i.e. 30 hpi.

We hypothesised that the increased intracellular ROS contribute to viral life cycle progression, which is in line with the observation that other viruses trigger ROS production to their own advantage (55). To test this hypothesis, we measured generation of new infective virions during the first virus cycle in cells treated with IKE, and with two strong antioxidant molecules authorised for clinical use, namely glutathione (GSH) and N-acetyl cysteine (NAC). In line with their antioxidant activity, GSH and NAC treatment reduced ROS levels (Fig. 5c). All treatments displayed strong antiviral activity (Fig. 5d) already after a single cycle of replication. These data suggest that the increase of ROS levels, especially at the early times post-infection, is instrumental for viral propagation. Thus, inhibiting this step may provide a new valuable therapeutic approach against SARS-CoV-2 infection.

While the manuscript was in preparation, the new variant of concern Omicron (BA.1) emerged and became dominant worldwide. In Fig. 5e we show that IKE and NAC treatments were also effective against Omicron indicating that our compounds target core processes shared among previous and emerging variants.

We then challenged the effects of the two candidate compounds in a mouse model of SARS-CoV-2 infection, namely K18-hACE2 mice. This model expresses the human ACE2 receptor under the promoter of keratin 18 in the epithelia, including the airway epithelial cells, and recapitulates several aspects of severe and non-severe COVID-19 in humans (56). Mice were treated daily with IKE, or NAC, for 4 days with one preinfection dose. Four days post-infection, the mouse lungs were harvested, total RNA purified, and viral load estimated by measuring the expression levels of two viral transcripts, i.e. the nucleocapsid (N) and the RNA dependent RNA polymerase (RdRp)(Fig. 6a). Remarkably, NAC-treated mice had a significantly lower expression of viral transcripts compared to untreated mice (Fig. 6b). We confirmed this result by immunohistochemical analysis of nucleocapsid protein in murine lungs (Fig. 6c and Supplementary Fig. 7). Thus, NAC is an effective treatment against SARS-CoV-2 infection in human cells and humanised mouse model of COVID-19.

**Figure 6.**
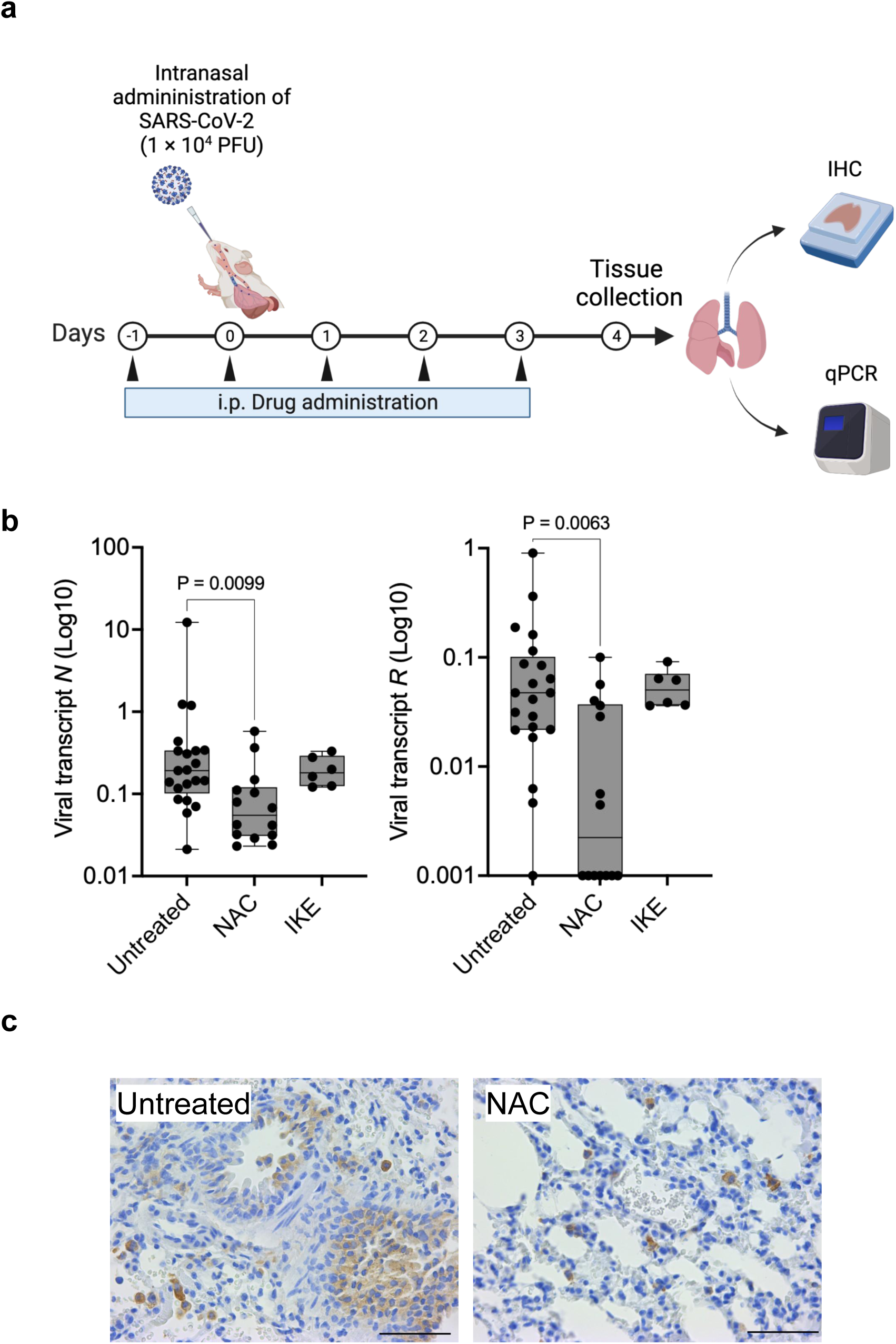
Treatment with NAC inhibits SARS-CoV-2 infection *in vivo*. **a**, Schematic of the experiment in vivo, created with Biorender.com. IHC, immunohistochemistry. **b**, qPCR analysis of viral transcripts (nucleocapsid, N; RNA dependent RNA polymerase, R) in lungs infected with SARS-CoV-2 and treated with NAC or IKE. n = 21 (Untreated), 14 (NAC), 6 (IKE) from 2 independent experiments. Data are presented as whisker plots: midline, median; box, 25-75th percentile; whisker, minimum to maximum values. Test: Kruskall-Wallis with corrected Dunn’s test for multiple comparisons. **c**, Representative images for viral nucleocapsid protein (N) stained by immunohistochemistry, for quantification see Supplementary Fig. 7. Scale bar: 100 μm.

## DISCUSSION

As obligate intracellular parasites, viruses tightly rely on their host cells: they have evolved to exploit cells for their own purposes by hijacking cellular pathways and to evade the innate immune response by modulating host factors and signalling pathways. RNA viruses, such as SARS-CoV-2, even more heavily rely on the host cell(57). However, current therapeutic interventions against COVID-19 are solely targeted against viral proteins, promoting the emergence of variants escaping vaccine-induced immunity or resistance to antiviral drugs.

The objective of this study was understanding whether targeting host proteins might be an effective and safe strategy against COVID-19, as host genes are not under selective pressure. We started by asking whether different SARS-CoV-2 variants elicit similar cellular responses upon infection. The three major SARS-CoV-2 variants of concern that we used in our study exhibited varying replication patterns in human lung cells, with Wuhan producing fewer transcripts and higher titres of infective viral progenies when compared to D614G and Alpha. We ascribed this diverse behaviour to the reported mutations in the Spike protein, which would result in modulated virus-receptor binding affinity and consequent viral entry(58). In addition, both D614G and Alpha harbour mutations and deletions in the open-reading frame 8 (ORF8) (Supplementary Table 1), which inhibits the cell interferon-mediated immune response(18, 59, 60), possibly explaining the enhanced RNA transcription of these two variants(61, 62). However, analyses of the host transcriptome revealed a qualitatively highly similar transcriptional response among the 3 variants, with differences in just kinetics and magnitude, overall indicating that different variants induce similar cellular responses upon infection.

We then applied a genome-wide CRISPR knockout approach to gather deep insights into the host genes exploited by different variants, asking whether some genes are specifically required for the infection of one or more variants. This kind of approach has been successfully developed to identify the host factors exploited by other viruses(63–67) and by SARS-CoV-2 itself(4–8, 68, 69). However, the analysis of available data of previous SARS-CoV-2 CRISPR knockout screenings does not allow us to draw conclusions about whether different variants exploit different host factors, because different studies used different combinations of variants, cell lines and CRISPR libraries. For these reasons, we performed a genetic screening directly comparing 3 variants under identical conditions and looked for the host factors that are conservatively exploited by all of them and, vice versa, those that are required by specific variants. The rationale of our approach is twofold: i) if a host factor is shared by all variants, it more likely belongs to a “core” of host factors essential for the viral infection and ii) shared host factors are more likely to be required by new variants of SARS-CoV-2 that will emerge in the future and thus might serve as a better and omni-comprehensive therapeutic target. By using conditions ensuring high coverage and stringency, we retrieved 525 genes, the knockout of which allowed cell survival upon infection; 93.3% were shared by at least 2 out of 3 variants. Very satisfactorily, all candidates selected by the CRISPR knock-out screening were also confirmed by transient silencing of host genes. Importantly, we failed to identify a single candidate acting specifically on only 1 variant. We conclude that the host factors exploited during infection are highly shared among different SARS-CoV-2 variants.

We believe that the knowledge acquired in this study will be instrumental to develop host-directed therapies to control SARS-CoV-2 infection. Due to their reliance on host cell components, these have reduced likelihood to develop resistance. To further assess the soundness of our hits and provide ready-to-trial drugs able to stop viral infection/replication of present and forthcoming variants, we screened a set of FDA-approved drugs against unrelated diseases, and chemical compounds reported to hamper the main common viral host factor candidates (SLC7A11, RIPK4 and MASTL). The five tested compounds displayed potent antiviral activity not only against the three tested SARS-CoV-2 variants, but also against the Delta variant, which appeared in late 2021 and has been so far the last variants that caused worrisome rates of hospitalisation of infected patients of all ages, regardless of their vaccination status, and was associated with high mortality rate(70, 71). The successful antiviral activity of the tested compounds further reinforces the strength of our screening, and points out that the selected hits are crucial host factors for both the early and latest variants.

The mechanism of action of one of the most promising tested compounds, IKE, was investigated to further validate its target, SLC7A11, against SARS-CoV-2. The central role of SLC7A11 in the maintenance of ROS intracellular homeostasis and its relevance as host factor in different human viral infections have been previously reported(37, 72–75). IKE was proposed to neutralise SLC7A11-mediated cystine uptake and ROS modulation(76). While increased intracellular ROS levels trigger innate immunity-mediated antiviral mechanisms, counterintuitively, viral infections stimulate ROS production and viruses thrive in increased ROS levels(55, 77). Indeed, our gene expression analysis suggests reduced oxidative phosphorylation within infected cells, possibly as an attempt the cells make to lower ROS and create a hostile environment for viral replication. We showed that SARS-CoV-2 stimulates ROS production during the early infective stages in human bronchial cells. Reduction of ROS levels, by extended IKE administration, glutathione or NAC treatment, impaired SARS-CoV-2 viral cycle. The effect of NAC treatment in COVID-19-affected patients has been investigated in several retrospective studies leading to suggestive, albeit not definitive, results (78–80). The mechanistic explanation was that the antioxidant, anti-inflammatory and anti-thrombotic effects of NAC counteracted viral pneumonia; however results from ongoing randomised controlled trials are required to draw accurate conclusions (79). In the meanwhile, our results show a direct antiviral effect of NAC on lung epithelial cells, in addition to its immunomodulatory effects. We thus strongly encourage and support NAC, and other antioxidant drugs, use as a safe and accessible anti-SARS-CoV-2 therapy, against current and future variants.

## MATERIALS AND METHODS

### Compounds

Tested compounds IKE (Cayman Chemical, US, Cat: 27088), MKI-1 (ChemBridge, US, Cat: 9335496), Sulfasalazine (MedChemExpress at MedChemTronica EU, Sweden, Cat: HY-14655), Tamatinib (Merck Life Science, Cat: 574714), Vandetanib (Merck Life Science, Cat: SLM2983) were dissolved in DMSO and stored in aliquots at −20°C until use. N-acetyl cysteine (NAC, Sigma-Aldrich, cat n. A9165) was reconstituted in sterile water and pH was adjusted to 7–7.4 with sodium hydroxide prior to use.

### Cell culture and virus

Vero E6 (ATCC® CRL-1586™) were maintained in Dulbecco’s modified Eagle’s medium (DMEM; Thermo Fisher Scientific), Calu-3 cells (ATCC®, HB-55) and Caco-2 cells (kind gift of Prof. Stefano Piccolo, University of Padua) were maintained in Dulbecco’s Modified Eagle Medium: Nutrient Mixture F-12 (DMEM/F-12, Thermo Fisher Scientific). Media were supplemented with 10% (v/v) fetal bovine serum (FBS, Thermo Fisher Scientific) and penicillin/streptavidin (Thermo Fisher Scientific). Cell cultures were maintained at 37°C and 5% CO_2_ in humidified atmosphere and routinely tested for mycoplasma contamination (Euroclone, Cat: EMK090020). For seeding and subcultivation, cells were first washed with phosphate buffered saline (PBS) and then incubated in the presence of trypsin/EDTA solution (Gibco, Thermo Fisher Scientific) until cells detached. Upon seeding of Calu-3 cells, 10 μM ROCK inhibitor Y27632 (Axon Medchem, Cat #1683) was added for 24 h to the culture medium.

The SARS-CoV-2 IT isolate SARS-CoV-2/human/ITA/CLIMVIB2/2020 was provided by the Virology Unit of Ospedale Luigi Sacco (GenBank accession ON062195 MW000351.1)(Milan, Italy). The SARS-CoV-2 USA isolate SARS-CoV-2/human/USA/USA-WA1/2020 was provided by The University of Texas Medical Branch (GenBank accession MT576563.1)(Galveston, USA). The SARS-CoV-2 UK and Delta isolates Human nCoV19 isolate/England/MIG457/2020 and hCoV-19/Netherlands/NH-RIVM-27142/2021_P2, respectively, were supplied by the European Virus Archive goes Global (EVAg) platform. The SARS-CoV-2 Omicron variant was provided by the Microbiology Unit of the University-Hospital of Padova (Padova, Italy), and previuosly described (GenBank accession ON062195) (81). All viral stocks were prepared by propagation in Vero E6 cells in DMEM supplemented with 2% FBS. Viral titre was assessed by plaque reduction assay (PRA) and expressed as plaque forming units (PFU) per milliliter (ml). All experiments involving live SARS-CoV-2 were performed in compliance with the Italian Ministry of Health guidelines for Biosafety Level 3 (BSL-3) containment procedures in the approved laboratories of the Molecular Medicine Department of University of Padova.

### Animal studies

Six- to 8-week-old B6.Cg-Tg(K18-ACE2)2Prlmn/J transgenic mice were purchased from The Jackson Laboratory and bred at the IOV-IRCCS Specific Pathogen-Free animal facility. K18-hACE2 mice were acclimatised in the BSL-3 facility for 72 hours prior to treatment. Mice were treated intraperitoneally either with 150 mg/kg of NAC or 40 mg/kg of IKE, or vehicles as controls. One day after the first dose (pre-treatment), mice were infected intranasally with 10 μl of 1 × 10^4^ PFU of SARS-CoV-2 Delta strain. Mice received drugs daily for 5 days, and were euthanized at day 4 post infection (dpi). Body weight and physiological conditions were monitored daily until sacrifice, when lungs were collected for further analysis. The whole right lobe was fixed in 10% buffered-formalin for histopathology, while the left lobes were added with Trizol (Invitrogen) for RNA extraction. All the procedures involving animals and their care were in conformity with institutional guidelines that comply with national and international laws and policies (D.L. 26/2014 and subsequent implementing circulars), and the experimental protocol (Authorization n. 355/2021-PR) was approved by the Italian Ministry of Health.

### SARS-CoV-2 titration by plaque reduction assay

Vero E6 cells were seeded in 24-well plates at a concentration of 9 × 10^4^ cells/well. The following day, serial dilutions of the viral stocks or tested supernatants were performed in serum-free DMEM media. After 1 h absorption at 37 °C, 2× overlay media was added to the inoculum to give a final concentration of 2% (v/v) FBS/DMEM media and 0.6% (v/v) methylcellulose (Merck Life science, Cat: M0512) to achieve a semi-solid overlay. Plaque assays were incubated at 37 °C for 48 h. Samples were fixed using 5% Formaldehyde in PBS (Merck Life Science, Cat: 252549) and plaques were visualised using Crystal Violet solution (20% Ethanol, Merck Life science, Cat: C6158).

### Virus infections in human lung cells and RNA sequencing analysis

Calu-3 cells were seeded (1.2×10^5^/well) in 12 well plate, and after 24 h the cell culture supernatant was removed and replaced with virus inoculum (MOI of 1 PFU/cell). Following 1 h adsorption at 37 °C, the virus inoculum was removed, the cell monolayer was washed in PBS prior to medium replacement (10% FBS DMEM/F-12). At 6, 9, 12 and 24 hours post infection (h.p.i.), cells were harvested and total RNA purified with Total RNA Purification Kit (NorgenBiotek, Canada, Cat #48400), following manufacturer’s protocol. Total RNA was retrotranscribed with random primer and M-MLV Reverse Transcriptase (Thermo Fisher Scientific, 28025013). qPCR analysis was carried out in a QuantStudio 6 Flex RealTime PCR System (Thermo Fisher Scientific) with FastStart SYBR Green Master mix (Roche, Cat. 04673492001). Primers for qPCR are listed in Supplementary Table 4.

### RNA sequencing

Total RNA was isolated with Total RNA purification kit (Norgen Biotek, Cat #48400) and Quant Seq 3’ mRNA-seq Library Prep kit (Lexogen) was used for library construction. Sequencing was performed on Illumina NextSeq 500 instrument with a coverage of ∼ 5 million reads (75bp SE) per sample. Raw reads obtained from RNAseq were mapped to a hybrid human (GRCh38.p13) and SARS-CoV-2 reference genome (GenBank: NC_045512.2) using STAR (v. 2.7.6a). The gene expression levels were quantified using Subread package featureCounts (v. 2.0.1). STAR parameters were set following Lexogen guidelines for data analysis (https://www.lexogen.com/quantseq-data-analysis/), while featureCounts was used with default parameters.

All RNA-seq analyses were carried out in the R environment (v. 4.1.0) with Bioconductor (v. 3.7). Genes were sorted removing those with a total number of raw counts below 10 in at least 4 samples. After applying this filter, we identified 11,880 expressed genes that were considered for further analyses. Outlier replicates were removed following quality control and clustering analysis. Differential expression analysis was computed using the DESeq2 R package (v. 1.32.0)(82). DESeq2 performs the estimation of size factors, the estimation of dispersion for each gene and fits a generalised linear model. Transcripts with absolute value of FC > 1.5 (log2[FC] > 0.59) and an adjusted p-value < 0.05 (Benjamini-Hochberg adjustment) were considered significant and defined as differentially expressed genes (DEGs) for the comparison in analysis. Volcano plots (Fig. 1b) were generated with log2[FC] and −log10[q-value] from DESeq2 differential expression analysis output using the ggscatter function from the ggpubr R package (v. 0.4.0). Heatmaps were made using DESeq2-normalised values with the pheatmap function from the pheatmap R package (v.1.0.12) on viral genes (Fig. 1a) or selected markers (Fig. 1c, 2d). Statistics and visualisation of the correlation matrix in Fig. 1d were performed with the Hmisc (v. 4.5-0) and Corrplot (v. 0.90) packages using Pearson’s correlation method.

Biological significance of DEGs was explored by GO term enrichment analysis (Fig. 1e, Supplementary Fig. 3 and Fig. 2b) using the enrichR package (v.3.0)(83).

### CRISPR-based loss-of-function screening

Calu-3 cells stably expressing SpCas9 were generated by transducing Calu-3 cells with lentivirus expressing Cas9 under the EFS promoter (provided with the library by Creative Biogene, see below) and selection with 2 μg/ml blasticidin. Transduction conditions were optimised to avoid non-specific effects of Cas9 on SARS-Cov-2 cytopathic effect. We conducted genome-wide negative selection (dropout) screens in Cas9-Calu-3 cells by using the human GeCKO v2 library (Creative Biogene, cat. CCLV0001) that targets 18823 genes with 6 gRNAs/gene as well as 1000 non-targeting gRNAs. The library is provided as two pooled DNA half-libraries (Library A and B, 3 gRNAs/gene each) that we screened in parallel. On day 1, four T175 flasks were seeded with 18,6×10^6^ Cas9-Calu-3 cells, the number of cells was optimised in order to have 15.5×10^6^ cells the following day. On day 2, 8.36 μl of Library A viral particles (stock 5.56×10^8^ TU/ml) and 7.92 μl of Library B viral particles (stock 5.87×10^8^ TU/ml) were resuspended in 20 ml medium and used to transduce two T175 flasks/semilibrary (20 ml/flask) with an MOI of 0.3. These conditions allowed a coverage of 500x, i.e. each gRNA is present, on average, in 500 unique cells and the majority of transduced cells received a single viral integrant. Transduced cells were selected with 4 μg/ml with puromycin and cultured for at least 2 weeks, after which cells transduced with the same semilibrary were pooled together. For each of the 4 conditions (infection with Wuhan, D614G and Alpha SARS-CoV-2, and a Mock sample) and each semilibrary, we plated 3 replicates, each of 6×10^6^ cells/replicate. These cells were spread into 12 wells (6-well format) to allow an even distribution of the cells (see Supplementary Fig. 5 for a schematics of the screening layout).

The day after, 72 wells (36 wells for each semilibrary) were infected, in parallel, with each of the SARS-CoV-2 variants at a MOI of 33. After 48 h, we observed complete death of non-infected control cells, and appearance of scattered clonal populations of cells that survived to SARS-CoV-2 infection. We expanded the colonies for 28 days in order to obtain a number of cells suitable for detection of each clone. The cell medium was changed every 48 hours. Then, all the wells transduced with the same gRNA library and infected with the same SARS-CoV-2 variant were lysed, pooled together and genomic DNA purified with phenol/chloroform. We considered this as a replica. Thus, for each gRNA semilibrary and SARS-CoV-2 variant we obtained and sequenced 3 replicates.

The gRNA cassettes of the surviving clones were identified as follows: gDNA from cells was extracted through Phenol-Chloroform and purified with ethanol precipitation to obtain the maximum extraction efficiency. The obtained gDNA was then purified using AmpureXP (Beckman A63881). Purified gDNA was quantified with Qubit 1x dsDNA High Sensitivity (Thermo Q33231) and subsequently used for PCR amplification. PCR was performed using KAPA HiFi HotStart ReadyMix (Roche #7958927001) at Tm 60°C for 15 cycles. The primers used were: GECKO2_Fwd: GCTTTATATATCTTGTGGAAAGGACGAAACACC; GECKO2_Rev: CCGACTCGGTGCCACTTTTTCAA. The PCR reaction was purified using Ampure XPbeads and run on 2% E-Gel™ EX Agarose Gels (Thermo G401002) to select a band of about 250 bp. DNA from agarose gel was purified using Zymoclean Gel DNA Recovery Kit (Zymo D4007). Obtained DNA was used for library preparation with NEBNext® Ultra™ DNA Library Prep Kit for Illumina® (NEB E7370L). Libraries were run on Novaseq 6000 (Illumina) on NovaSeq 6000 SP Reagent Kit v1.5 (100 cycles) (Illumina 20028401).

Data processing was conducted using the MAGeCK software(84) in combination with a custom pipeline. Briefly, read counts from different samples were first mapped to the reference gRNA sequences library using “mageck count” function with default parameters; as the sequencing library is unstranded, reads were mapped also to the reverse complement of the gRNA library and then counts were combined.

Individual gRNA-level and aggregate gene-level enrichment analysis was performed using a custom pipeline. gRNA counts from different samples were normalised to total counts to adjust for the effect of library sizes. Only gRNAs with a count number higher than the maximum count value of control samples (CTRL, cell transduced with the GeCKO library that were not infected) were considered enriched and thus retained for further analyses.

We calculated a gRNA score that represents the number of biological replicates in which a gRNA for a given gene were found enriched over control samples. The gRNA score was calculated within each variant (ranging from 0 to 3 replicates) or combining all variants (ranging from 0 to 9 replicated).

Genes were considered screen hits if targeted by at least 2 independent gRNAs and if the number of counts > 1000 at least in one sample; to increase stringency of the analysis, we considered only genes with a total gRNA score > 2 and we also calculated the average expression in Calu-3 cells and filtered out genes with <30 normalised counts.

### Protein network analysis

Protein network was generated with STRING 11.5 (online tool: https://string-db.org/) and Cytoscape stringApp. A first network was generated with STRING by using all the proteins belonging to the following libraries: Wikipathways (Host-pathogen interaction of human coronaviruses - interferon induction WP4880, Type I interferon induction and signalling during SARS-CoV-2 infection WP4868), GO Biological Processes (mitochondrion organisation (GO:0007005), negative regulation of TOR signalling (GO:0032007)), GO Molecular Functions (ribosomal large subunit binding (GO:0043023), RNA binding (GO:0003723), oxidoreductase activity, acting on the aldehyde or oxo group of donors, NAD or NADP as acceptor (GO:0016620)). This network was exported to Cytoscape and manually curated to facilitate visualisation by removing unconnected nodes (score > 0.70). Edges show connections based on experiments, coexpression, text mining, databases, cooccurrence, neighbourhood, fusion (any) and edges width is proportional to the strength of the interaction (mapping type “continuous” based on “Stringdb score” value).

### Validation of candidate genes

Selected candidate genes were validated in Calu-3 and Caco-2 cell lines by transient transfection. Cell reverse transfections were carried out using HiPerFect (Qiagen, 301704) for Calu-3 (2.5×10^4^ cells/well in 96well format) and forward transfections with Lipo3000 were done (Thermofisher, L3000015) for Caco-2 cells (1×10^4^ cells/well in 96well format). The siRNAs were selected from the FlexiTube GeneSolution 4 siRNA sets (Qiagen) and transfected as a mix at 24 nM in Calu-3 and 10 nM in Caco-2 following manufacturer’s instructions. As a negative control for our transfections we used a non-targeting siRNA from Qiagen (SI03650318, sequence: UUCUCCGAACGUGUCACGU). Cells were harvested 48 h post-transfection, their total RNA was purified and retrotranscribed as in Ref.(85). Real-time PCR was performed as in Ref.(86) with primers listed in Supplementary Table 4.

At 24 h post-transfection, the cell culture supernatant was removed and replaced with virus inoculum (MOI of 0.1). Following 1 h adsorption at 37 °C, the virus inoculum was removed and replaced with fresh 10% FBS DMEM/F-12 media. Cells were incubated at 37 °C for 48 h before supernatants were harvested. The viral titre (expressed as PFU/ml) was calculated by PRA in Vero E6 cells.

### Cytotoxicity evaluation of tested compounds

The cytotoxicity of the tested compounds was assessed and expressed as cytotoxic concentration (CC_50_). Calu-3 cells (2.75×10^4^ cells/well) were seeded in 96 well plates and the tested drugs or an equal volume of vehicle (DMSO) were supplemented to the medium. Compounds were incubated for 48 h and cell viability was determined by measuring the adenosine triphosphate (ATP) content of the cells using the ATPlite kit (PerkinElmer, Waltham, MA, Cat: 6016941) according to the manufacturer’s instructions. CC_50_ values were calculated using the Reed and Muench method(87).

### Antiviral assays

Calu-3 cells (2.75×10^4^ cells/well) were seeded in 96 well plates and the tested compounds or an equal volume of vehicle (DMSO) were supplemented to the medium 24 h prior to infection. The cell culture medium was removed and replaced with virus inoculum (MOI of 0.1 PFU/cell). Following 1 h adsorption at 37 °C, the virus inoculum was removed and replaced with fresh 10% FBS DMEM/F-12 media supplemented with the tested compounds or the vehicle. Cells were incubated at 37 °C for 48 h before supernatants were harvested. The viral titre (expressed as PFU/ml) was calculated by PRA in Vero E6 cells. IC_50_ values were calculated using the Reed and Muench method(87).

### ROS measurements

ROS measurement was performed by H2DCFDA assay, according to the manufacturer’s instructions (Thermo Fisher Scientific, D399). In brief, Calu-3 cells (2.75×10^4^ cells/well) were seeded in 96 well plates. Tested compounds or an equal volume of vehicle (DMSO) were supplemented to the medium 24 h prior to ROS analysis, if not otherwise stated. Each condition was tested in sextuplicate. Following drug treatment, media was removed and cells were incubated with 10 μM H2DCFDA in phenol-red free media for 20 min at 37 °C. Cells were washed with clear medium to remove free probe and fluorescence intensity (excitation=485 nm; emission=530 nm) was measured using a microtiter plate reader (Promega, GloMax Microplate reader). In each experimental plate, an additional lane of control cells was treated for 3 min at 37°C with H2O2 (3.6% w/v), to test probe correct fluorescence.

### Assessment of viral transcripts *in vivo*

At 4 days post infection, lungs were harvested in 2 ml of Trizol and homogenized using a gentleMACS Octo dissociator (Miltenyi Biotec, Inc.). Total RNA was purified with trizol/chloroform, genomic DNA digested with DNAse (DNAse I, Ambion, Thermofisher AM222) treatment followed by a second round of purification with phenol/chloroform/isoamyl alcohol. qPCR analysis was carried out in a QuantStudio 6 Flex RealTime PCR System (Thermo Fisher Scientific) with TaqPat 1-Step RT-qPCR (Applied Biosystems, Thermofisher, A15299). Primers for qPCR are listed in Supplementary Table 4. Probe for N transcript: CTAACAATGCTGCAATCGTGC (Reporter: FAM; Quencher: TAMRA). Probe for R transcript: CTATATGTTAAACCAGGTGGAACC (Reporter: FAM; Quencher: TAMRA). Probe for *ApoB* transcript: CCA ATG GTC GGG CAC

TGC TCA A (Reporter: VIC; Quencher: TAMRA). Expression of viral transcripts in each sample was calculated with the formula 2^(Ct *ApoB*-Ct N/R). Statistical analysis was performed with Graphpad Prism 9 Version 9.4.1 (458).

### Immunohistochemistry of murine lung tissue

After tissue harvesting, the right lung lobe was fixed in 10% buffered-formalin, dehydrated through a graded series of ethanol and embedded in paraffin (FFPE). Immunohistochemical (IHC) examinations of 4μm thick lung sections were performed on polarised glass slides (TOMO, Matsunami Glass IND, Osaka). Heat-induced antigen retrieval with 0,01 M Sodium citrate buffer, pH 6.0 for 60 minutes, at 97°C was followed by blocking of nonspecific bindings with 5% bovine serum albumin. Primary anti-SARS-CoV-2 nucleocapsid rabbit polyclonal antibody (Pro Sci Incorporated, Flint, CA, Cat: 9099, 1:300) was applied overnight at room temperature in a humidified chamber. Slides were then incubated with a HRP-conjugated secondary anti-rabbit antibody (Invitrogen, Carlsbad, CA, Cat: 31460, 1:500) for 60 minutes at room temperature. After endogenous peroxidase blocking (Agilent Technologies, Santa Clara, CA, Cat: S2023), 3,3’-diaminobenzidine peroxidase substrate detection kit (Agilent Technologies, Santa Clara, CA, Cat: K3467) was used to detect immunoreactivity. Non-infected murine pulmonary tissue was used for negative controls. Intensity of signal was subjectively scored in different anatomical compartments (i.e. blood vessels, interstitium, an airways/alveoli) as follows: 0, not detected; 1, mild/weak; 2, moderate; 3, strong. Nucleocapsid protein is detected primarily in alveolar pneumocytes type II and interstitial macrophages. Finally, a total IHC cumulative score for each section was obtained.

## Code availability

All custom code generated for RNAseq and CRISPR screening analyses is available upon request.

## Data availability

Raw sequencing data (RNAseq and CRISPR screening) have been deposited on GEO (accession: GSE207981). Source data are provided with this paper.

## ACKNOWLEDGEMENTS AND FUNDING

We thank Sirio Dupont (University of Padua), Milena Bellin (University of Padua) and Lucia Nencioni (Sapienza University, Rome) for critical reading of this manuscript. We thank Sirio Dupont and Patrizia Romani for providing reagents and suggestions on experiments related to ROS. We also thank Fenderico Manuel Giorgi for help with transcriptomic analysis. This work was supported by grants from the CaRiPaRo foundation titled “Identification of novel antiviral drug targets by genome-wide screening of cellular genes crucial for SARS-CoV-2 infection” to G.M. and “Interazione virus ospite nella pandemia COVID-19: studio immuno-virologico per la comprensione della patogenesi e la individuazione di bersagli terapeutici in una popolazione “fragile” di pazienti oncologici” to A.R. and from the Ministry of Education, University and Research “Dissecting the complex network of virus-cell host interactions controlling viral replication and inflammatory response to identify novel host-targeted approaches against severe respiratory virus infections (INHALA)” (PRIN-2020 2020KSY3KL) to S.N.R.. The Martello lab is supported by grants from the Giovanni Armenise–Harvard Foundation, the Telethon Foundation (TCP13013) and an ERC Starting Grant (MetEpiStem). The Richter lab is further supported by grants from the European Research Council (ERC Consolidator 615879), the Bill and Melinda Gates Foundation (OPP1035881 and OPP1097238), the Italian Foundation for Cancer Research (AIRC 21850) and the Collaborative Center for XDP at Massachusetts General Hospital (239295). The Montagner lab is supported by AIRC under MFAG 2021 (ID 25745 project) and “STARS Consolidator Grant” from University of Padua. The Rosato lab is supported by Fondazione AIRC-IG 2018 (ID 2135), Italian Health Ministry’s RCR-2019 (ID 23669115) and NET-2016 (ID 02361632), Ricerca Corrente funding from the Italian Ministry of Health and Veneto Institute of Oncology IOV-IRCCS. This work was supported by Fondazione Telethon Core Grant, Armenise-Harvard Foundation Career Development Award, European Research Council (grant agreement 759154, CellKarma) to DC, and grants from Regione Campania (G84I2000330005,G85F21000040002) to TIGEM.

This project was supported by the European VirusArchive GLOBAL (EVA-GLOBAL) project that has received funding from the European Union’s Horizon 2020 research and innovation programme under grant agreement No 871029, which provided access to SARS-CoV-2 UK and Delta isolates (Human nCoV19 isolate/England/MIG457/2020 and hCoV-19/Netherlands/NH-RIVM-27142/2021_P2).

## Author contributions

S.N.R., M.M. and G.M. conceived the study. I.F., C.S., A.D.P and A.P. performed BSL3 experiments. L.D. performed computational analyses. E.C. performed cell culture and molecular biology experiments. L.V. and D.C. performed NGS on CRISPR screening samples. A.D.P performed experiments with humanised mouse models. R.V. and F.T. performed histopathological analyses. A.R., S.N.R., M.M. and G.M. provided overall supervision and secured grants. M.M., L.D. and I.F. prepared figures. I.F., S.N.R., M.M. and G.M. wrote the manuscript with help from all the authors.

---

**Supplementary Figure 1.**
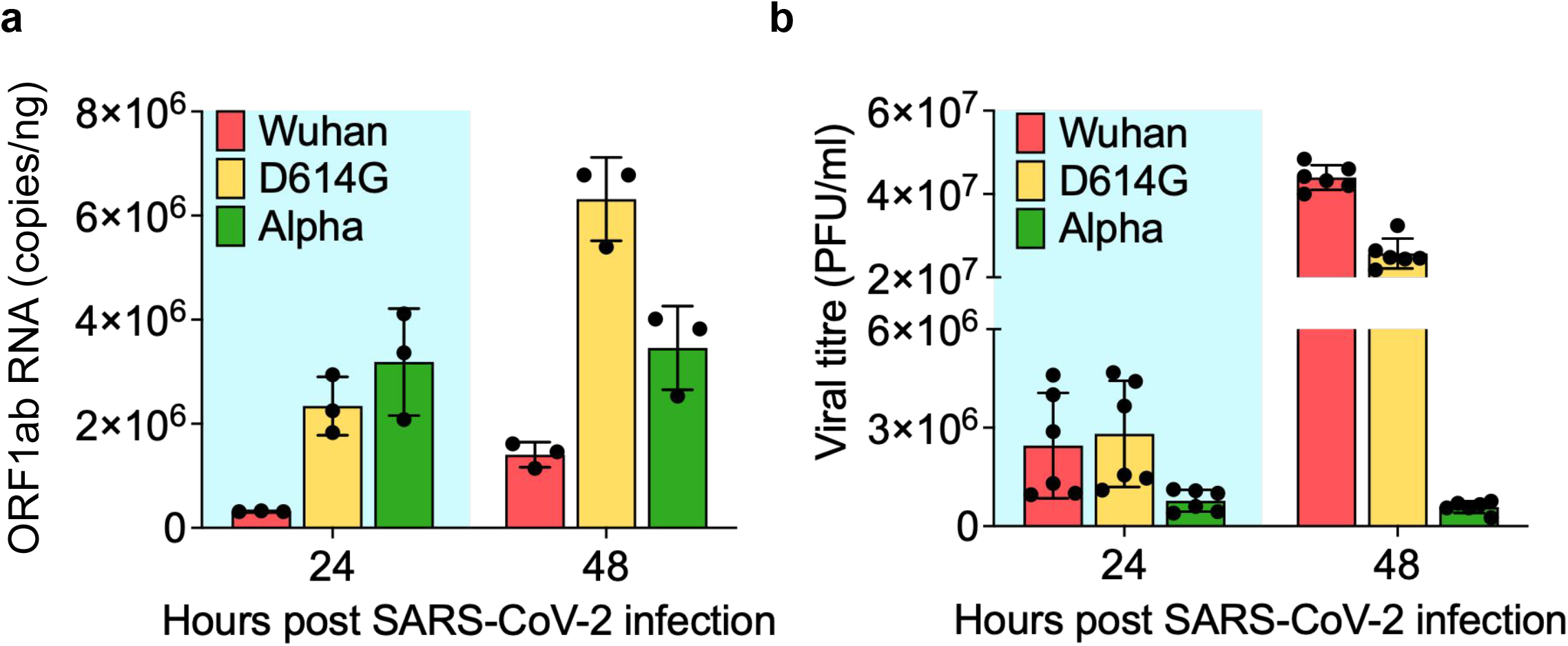
SARS-CoV-2 variants’ replication kinetics. Calu-3 cells were infected with the SARS-CoV-2 Wuhan, D614G, and Alpha variants (MOI 0.1). The viral intracellular RNA (**a**) and the viral particles released in the supernatant (**b**) were measured over time (24 and 48 h.p.i.). The RNA copies of the ORF1ab gene, specifically the region encoding for the RNA-dependent RNA polymerase, were calculated using a standard curve generated with a Rpdp-amplicon encoding plasmid. The viral titre was calculated by plaque reduction assay and expressed as PFU/ml. Data are mean ± s.d. of n=3 biological replicates in **a**, and of n=2 biological replicates, each tested in technical triplicate in **b**.

**Supplementary Figure 2.**
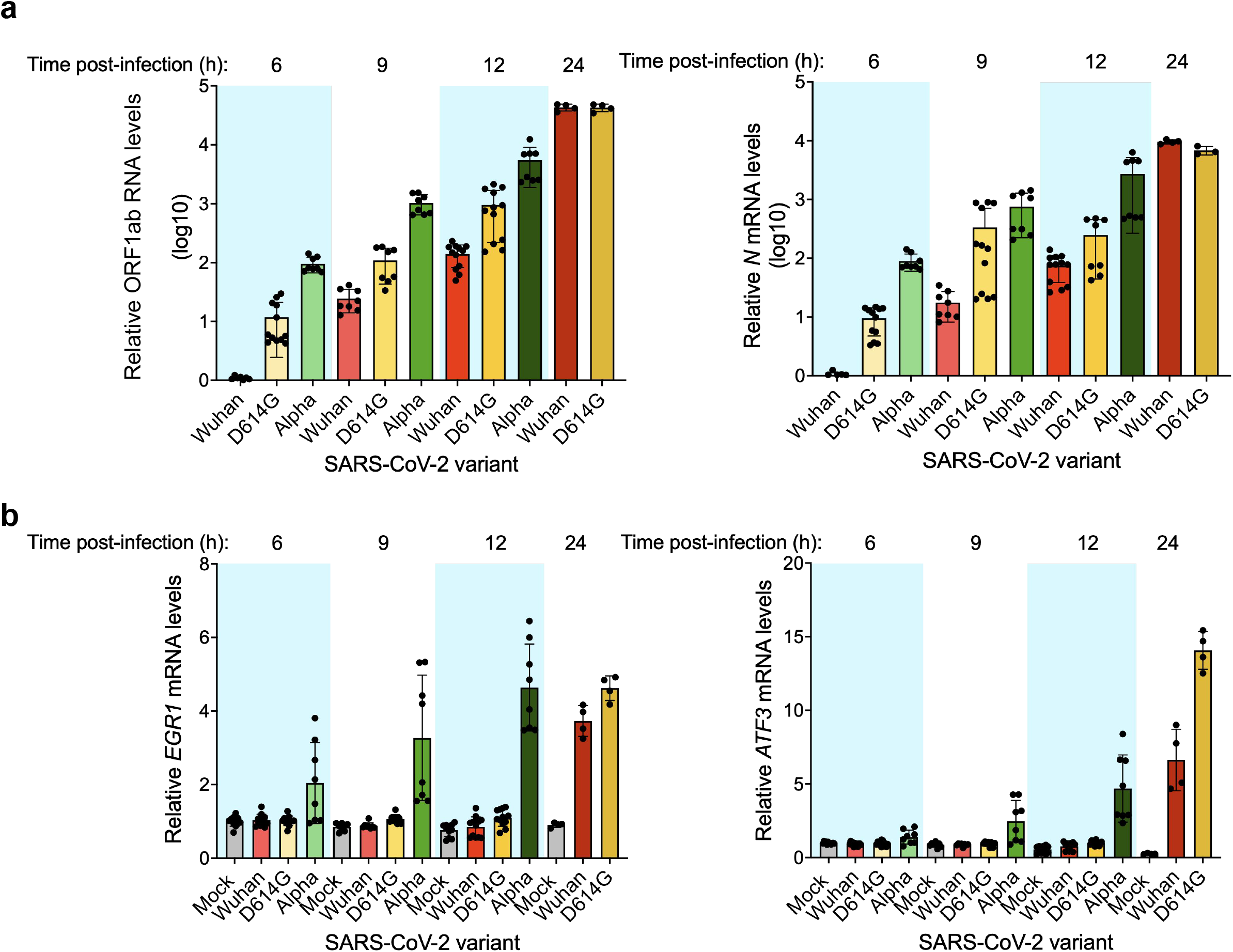
qPCR validation of viral and cellular transcripts. Relative expression of (**a**) viral genes (*ORF1ab* and *N*) and (**b**) cellular genes (*EGR1* and *ATF3*) in Calu-3 cells infected with the SARS-CoV-2 Wuhan, D614G, and Alpha variants (MOI 1). Intracellular RNA levels were measured at 6, 9, 12, 24 h.p.i. by qPCR. Data are mean-normalised pooled values from independent experiments (n=2 for Alpha; n=3 for Wuhan and D614G variants). Each condition was tested in 4 replicates per condition in each experiment. EGR1 and ATF3 are genes previously identified by Wyler and colleagues^22^ as strongly induced at 12 h.p.i.

**Supplementary Figure 3.**
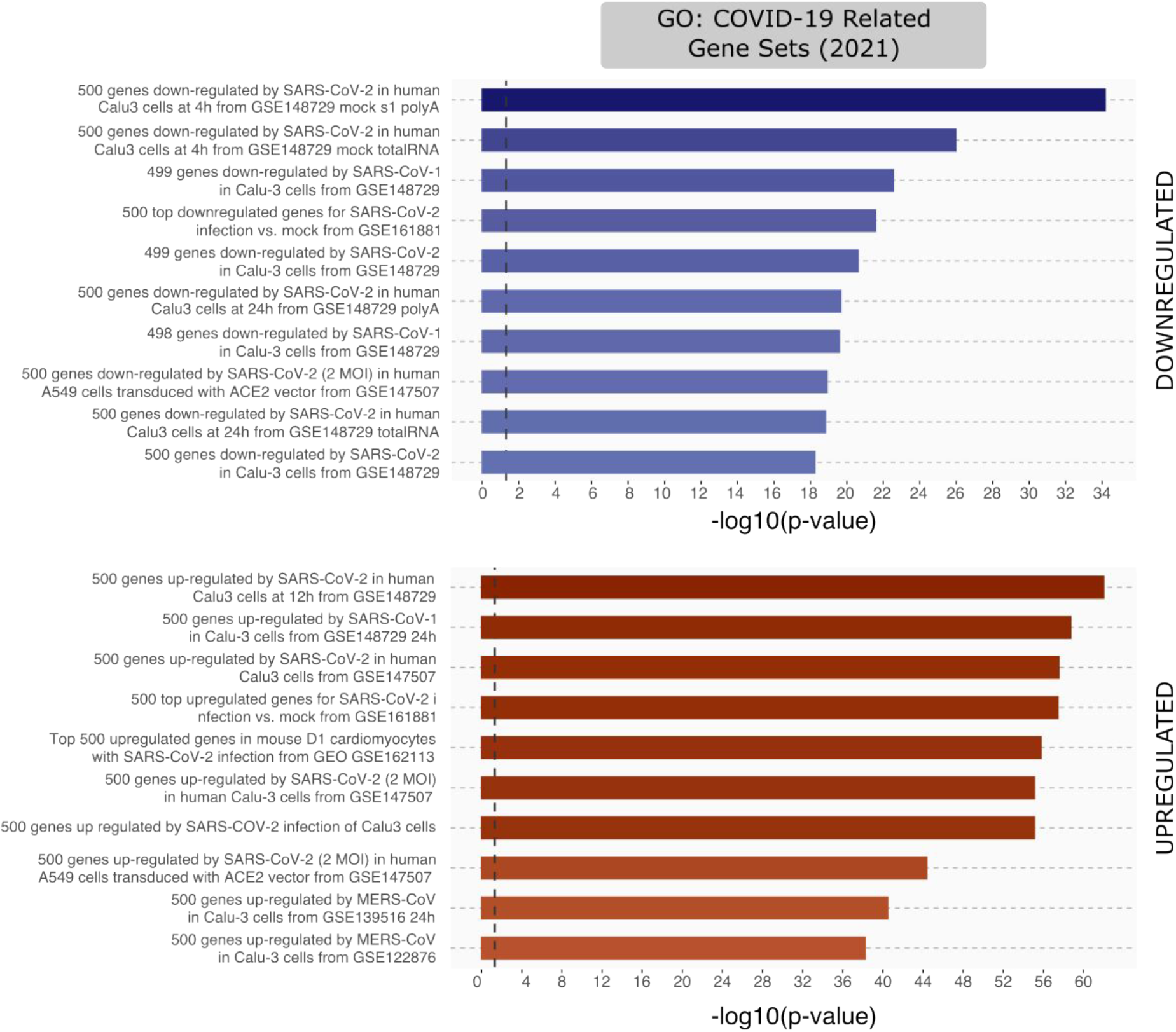
Gene enrichment analysis on DEGs identified in samples infected with Alpha variant for 12 h.

**Supplementary Figure 4.**
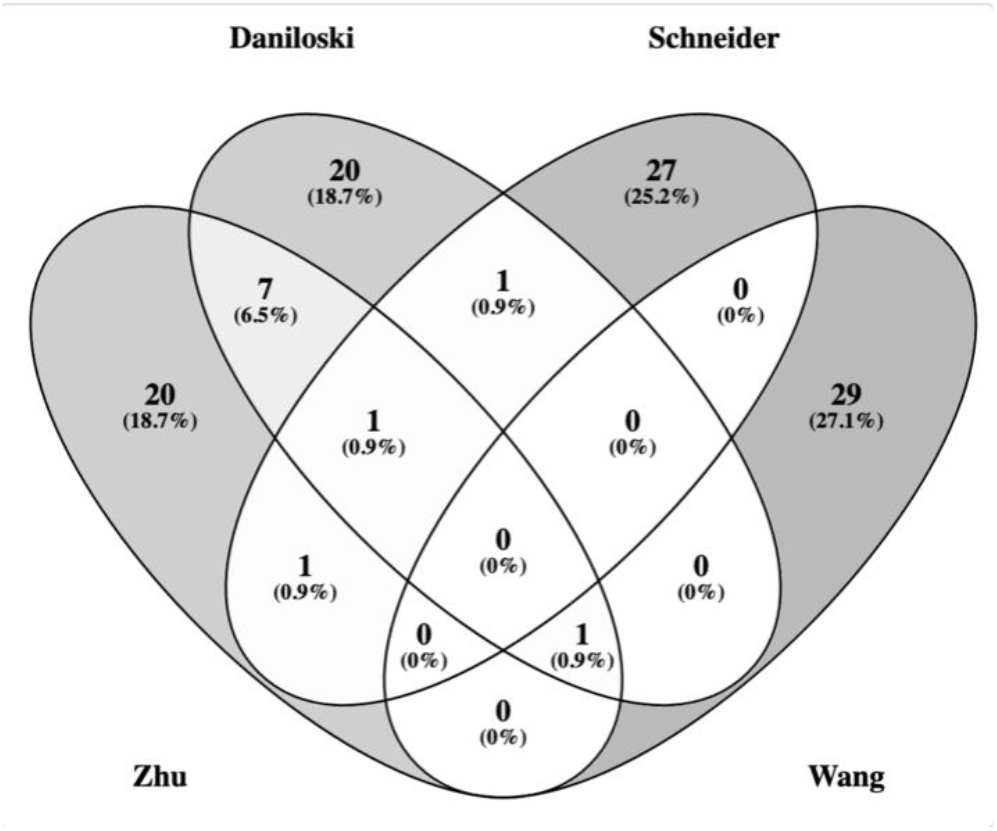
Overlap of cellular genes identified in the different CRISPR-based genetic screens.

**Supplementary Figure 5.**
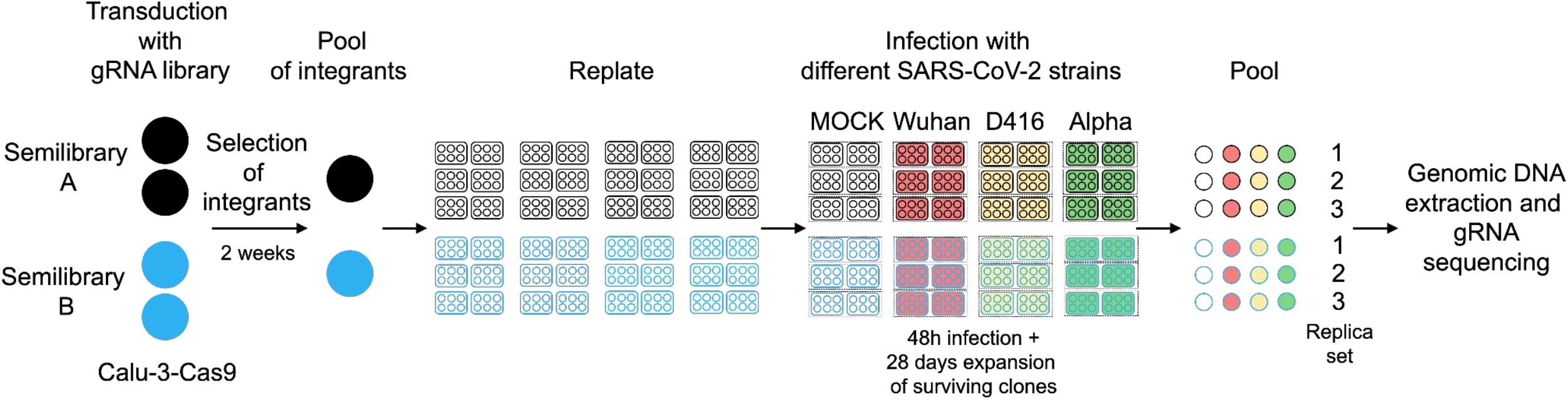
Workflow of CRISPR-based loss-of-function screening.

**Supplementary Figure 6.**
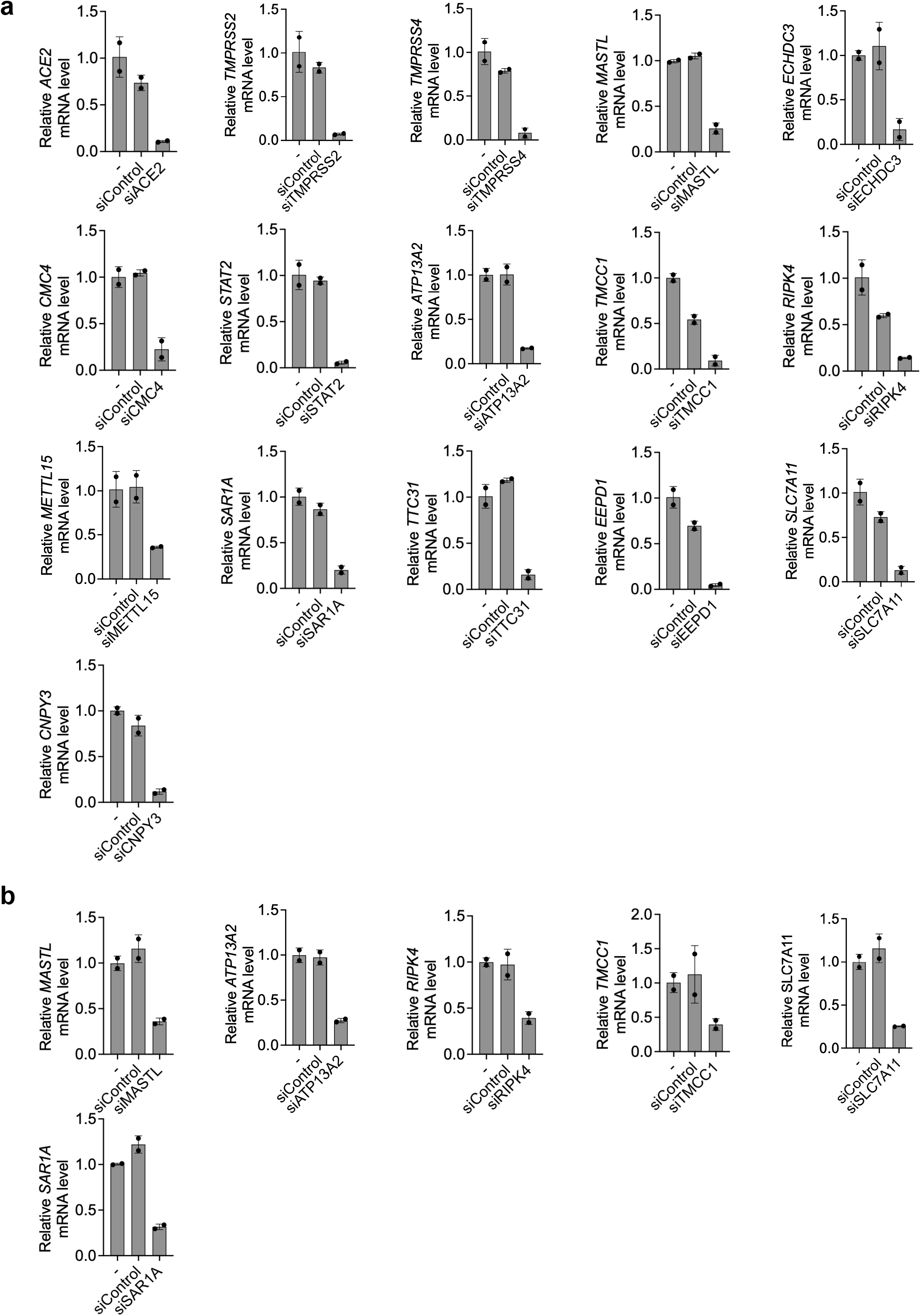
siRNA-mediated knock-down efficiency in human cells. Relative expression of selected candidate hits after siRNA knockdown in (**a**) Calu-3 and (**b**) Caco-2 cell lines. RNA levels were measured by qPCR 48 hours post siRNA transfection. Data are mean ± s.d. of n=2 biological replicates.

**Supplementary Figure 7.**
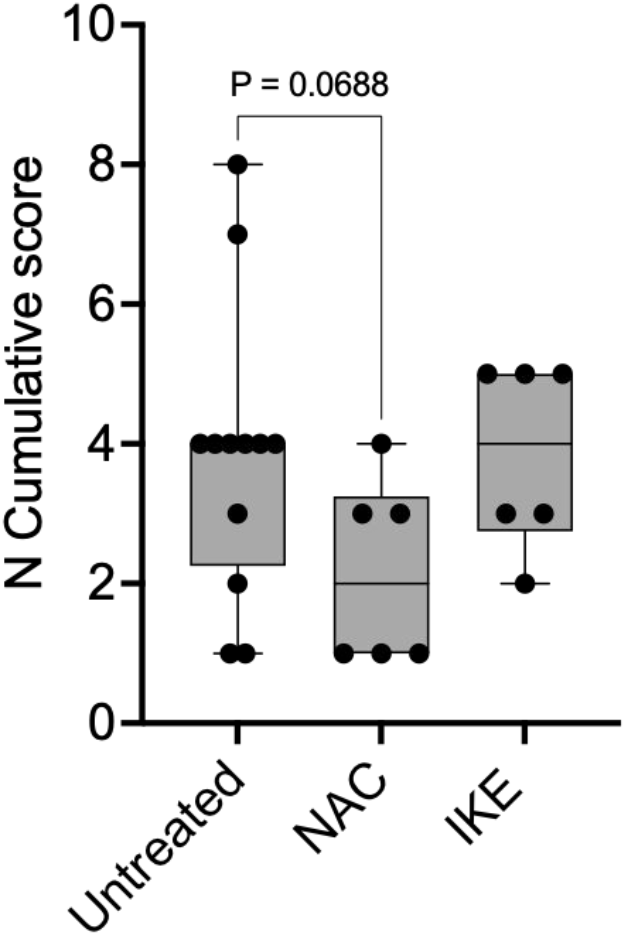
Immunohistochemical analysis of SARS-Cov-2 nucleocapsid protein in humanised model of SARS-CoV-2 infection treated with NAC and IKE. N cumulative score has been calculated as described in Methods section and representative images are shown in Fig. 6c. Data are presented as whisker plots: midline, median; box, 25-75th percentile; whisker, minimum to maximum values. Test: Kruskal-Wallis with uncorrected Dunn’s for multiple comparisons.

**Supplementary Table 1.** Amino acidic mutations present in the SARS-CoV-2 variants used in this study. Only mutated positions are reported. The Wuhan variant was used as reference strain, as it represented the first circulating SARS-CoV-2 isolate. Each mutation (such as A1707D) is indicated by a first letter that is the symbol for the reference amino acid on the reference Wuhan variant (e.g. A), a number for the amino acid position in the analyzed variant, and a second letter representing the amino acid actually found in the analyzed sequence (e.g., D).

| Protein/strain | Wuhan | D614G | Apha | Delta |
| --- | --- | --- | --- | --- |
| ORF1AB (7096 aa)<br>*same aa residue |  |  | A1707D<br>A2123V<br>Del 3673-3675 LSG<br>F3676*L<br>N4355K<br>K5781R<br>E6282G | Del 141-143 KSF<br>P306*L (P309)<br>P1637*L (P1640)<br>D2878*N<br>F3135*S<br>H3577*Q<br>K6708R |
| Orf1ab leader prot (nsp1) |  |  |  | Del 141-143 KSF |
| nsp2 |  |  |  |  |
| nsp3 (IT from MT seq) |  |  |  |  |
| nsp4 (IT from MT seq) |  |  |  | D217N<br>F375S |
| nsp3C-like protease |  |  |  |  |
| nsp6 |  |  | Del 102-105 LSG<br>F105L | H11Q |
| nsp7 |  |  |  |  |
| Nsp8 |  |  |  |  |
| Nsp9 |  |  |  |  |
| Nsp10 |  |  |  |  |
| Nsp11 (not considered a<br>viral protein any longer) |  |  |  |  |
| RNAdeprNApol (nsp12) |  | L623P |  |  |
| Helicase (nsp13) |  |  | K460R |  |
| 3'-5' Exonuclease (nsp14) |  |  | E347G |  |
| Endoribonuclease (nsp15) |  |  |  | K259R |
| 2'-O-ribose<br>methyltransferase (nsp16) |  |  |  |  |
| S |  | D614G | Del69-70 HV<br>Del145Y<br>N498Y<br>A567D<br>P678H<br>T713I<br>D1115H | T19R<br>K77T<br>G142D<br>Del156-57 EF<br>L452R<br>T476K<br>P679R |
| E |  |  |  |  |
| M |  |  |  | I82T |
| ORF6 |  |  |  |  |
| ORF7a |  |  |  | V82A |
| ORF7b |  |  |  |  |
| ORF8 |  | L84S | Truncated at 26th AA<br>(nt 27956 stop codon) |  |
| N |  |  | D3L<br>R203K<br>G204R | D63G<br>R203M<br>S235F<br>D377Y |
| ORF10 |  |  |  |  |

**Supplementary Table 3.**
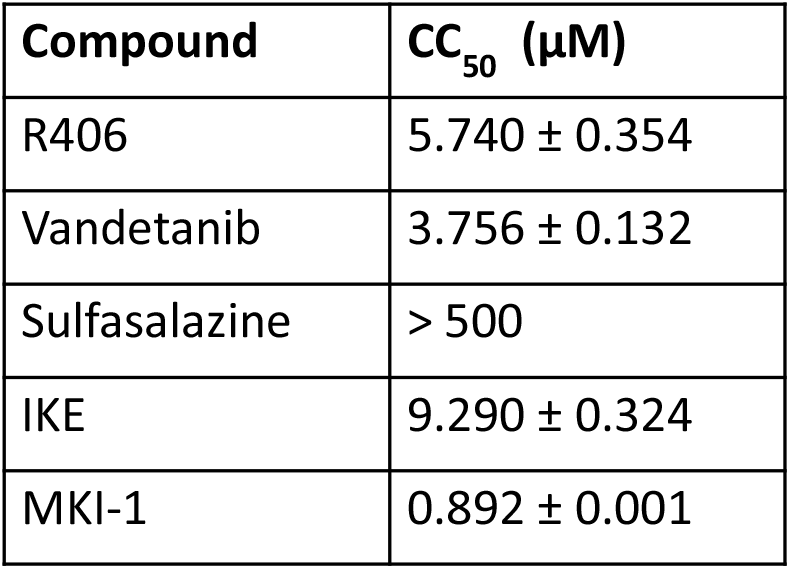
The cytotoxicity (CC50) of the the used drugs was determined in Calu-3 cells. Data are mean ± s.d. of n = 2 biological replicates. Each biological replicate included three technical replicates.

**Supplementary Table 4:**
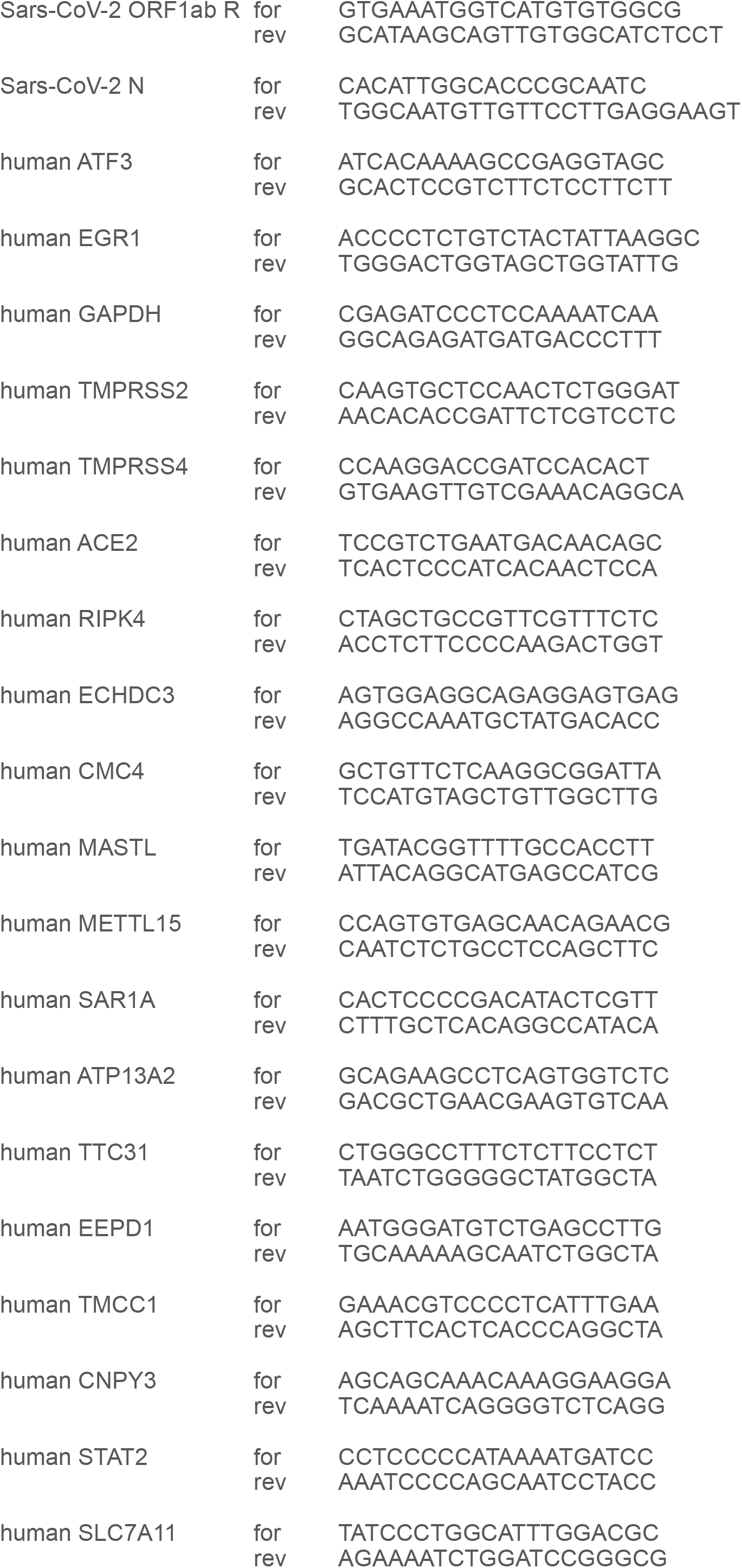
Sequence of primers used in this study.

**Supplementary Table 5:**
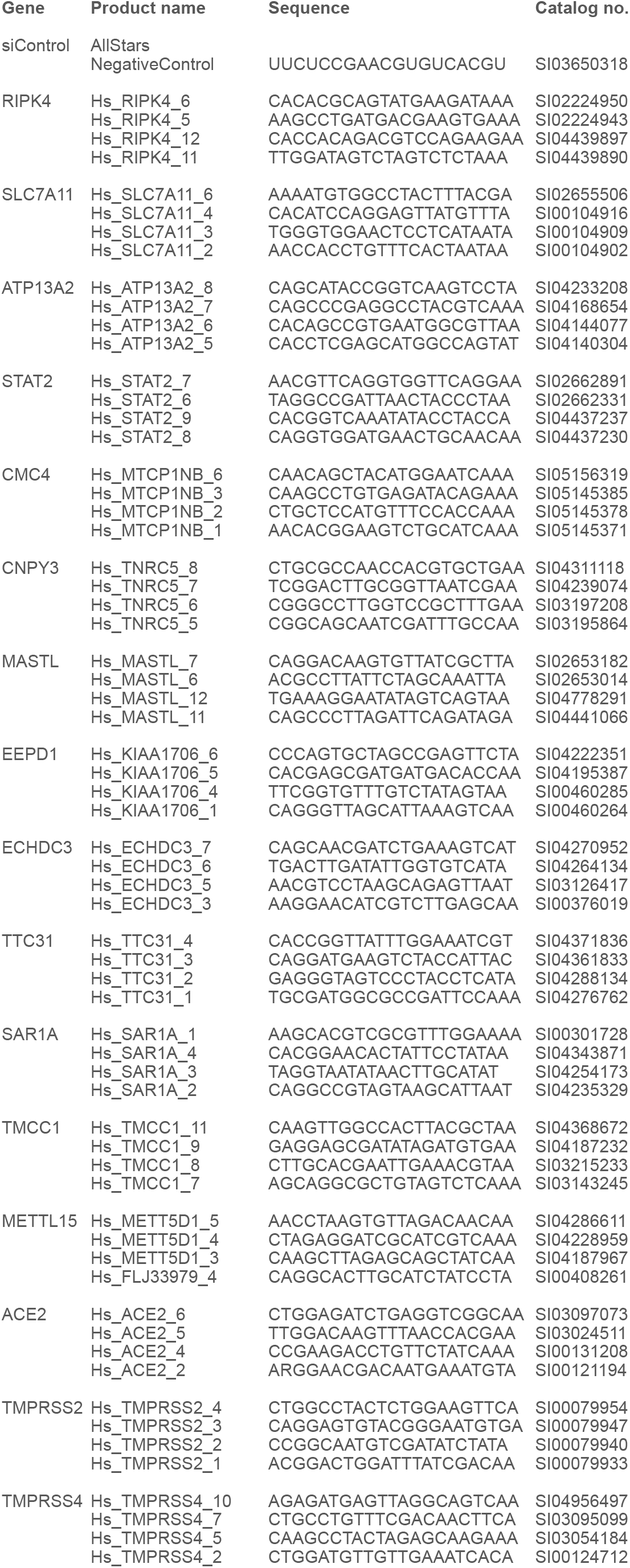
Sequence of siRNA used in this study.

